# Self-assembly of mycolic acid in water: monolayer or bilayer

**DOI:** 10.1101/2024.08.13.607716

**Authors:** Yogendra Kumar, Subhadip Basu, Prabal Kumar Maiti

## Abstract

The enduring pathogenicity of Mycobacterium tuberculosis can be attributed to its lipid-rich cell wall, with mycolic acids (MAs) being a significant constituent. Different MAs’ fluidity and structural adaptability within the bacterial cell envelope significantly influence their physicochemical properties, operational capabilities, and pathogenic potential. Therefore, an accurate conformational representation of various MAs in aqueous media can provide insights into their potential role within the intricate structure of the bacterial cell wall. We have carried out the MD simulations of MAs in an aqueous solution and shed light on the various structural properties such as thickness, order parameters, area-per-MAs, conformational changes, and principle component (PC) in the single component and mixed MAs monolayer. The different conformational populations in the monolayer have been estimated using the distance-based analysis between the function groups represented as W, U, and Z-conformations that lead to the fold of the MAs chain in the monolayer. Additionally, we have also simulated the mixture of alpha-MA (α-MA or AMA), methoxy-MA (MMA), and keto-MA (KMA) with 56% AMA, 40 % MMA, and 14 % KMA composition. The thickness of the MAs monolayer was observed to range from 5 nm to 7 nm with an average 820 kg/m^3^ density for α-MA, MMA, and KMA quantitative agreement with experimental results. The mero-chain (long chain), consisting of a functional group at the proximal and distal positions, tends to fold and exhibit a more disordered phase than the short chain. The keto-MA showed the greatest WUZ total conformations (35.32 %) with decreasing trend of eZ > eU > aU > aZ folds in both single component and mixture. Our results are in quantitative agreement with the experimental observations. The most minor population of sZ folds (0.75 % in a single component and 1.1 % in a mixture). However, eU and aU folds are most probable for the AMA and MMA. One striking observation is the abundance of MA conformers beyond the known WUZ convention because of the wide range distribution of intramolecular distances and change in dihedral angles. From the thermodynamic viewpoint, all monolayers were found to be stable in nature. The results from enhanced simulation revealed that changes in the magnitude of minimum free energy of the MAs are in order of KMA > AMA > MMA, and keto-MA exhibits more resistance (based on free energy values) for the drug molecules to permeate through the monolayer.

## 1. Introduction

The worldwide tuberculosis (TB) pandemic poses a significant risk to human well-being and progress, necessitating immediate intervention. Approximately 90% of the annual incidence of TB is observed in the adult population, with a higher prevalence among males than females.^1^ In 2021, the recorded global mortality rate attributed to TB was 1.4 million deaths.^1^ Furthermore, the impact of the COVID-19 pandemic on mortality from TB has been significantly more significant than that on HIV/AIDS mortality.^1^

The genus Bacillus encompasses a diverse group of bacteria^2^, including *Mycobacterium*. *Mycobacterium tuberculosis* (*M. tb.*) serves as the etiological agent responsible for the development of tuberculosis (TB) infection in the human population. In general, the treatment of mycobacterial infections poses challenges due to the inherent resistance of mycobacteria to a wide range of commonly used antibiotics and chemotherapeutic drugs.^3^ One of the contributing aspects to the development of resistance is the impediment posed by the cell wall of mycobacteria.^4–9^ The mycobacterial cell wall consists of many layers and exhibits exceptional thickness and impermeability. The layer of *m.tb* cell wall that is of particular interest due to its permeability property is known as mycomembrane.^10^ The outer leaflet of the mycomembrane is comprised of free lipids like trehalose dimycolate (TDM), Trehalose Monomycolate (TMM), and various other glyco and phospholipids.^10^ On the other hand, a tightly packed monolayer of mycolic acid (MA), covalently bonded with arabinogalactan (AG), a polysaccharide, forms the inner leaflet of mycomembrane. The Mycomembrane and specifically the mycolic acid layer within it is speculated to be responsible for the drug resistance property of the bacteria, as it acts as a highly effective permeability barrier.

Mycolic acids (MAs) are 2-alkyl, 3-hydroxy long fatty acids (FAs) with distinctive properties (as shown in Figure 1).^11^ The initial structural elucidation of MAs occurred in 1950, wherein they were identified as long-chain FAs possessing two branches and three hydroxyl groups.^12^ The functions of MAs exhibit a quantifiable influence on various aspects, including cell wall permeability, growth, virulence, etc.^13–18^ MAs exhibit a considerable range of chain lengths and chemical functionalities, which are responsible for distinguishing several classes of mycolic acids.^18,19^ It has been speculated that a recent study employed a machine learning approach to develop a neural network model for predicting the permeability of drug-like molecules through M. *tb* cell wall, which will be useful to identify potential drug candidates. ^20^

**Figure 1.**
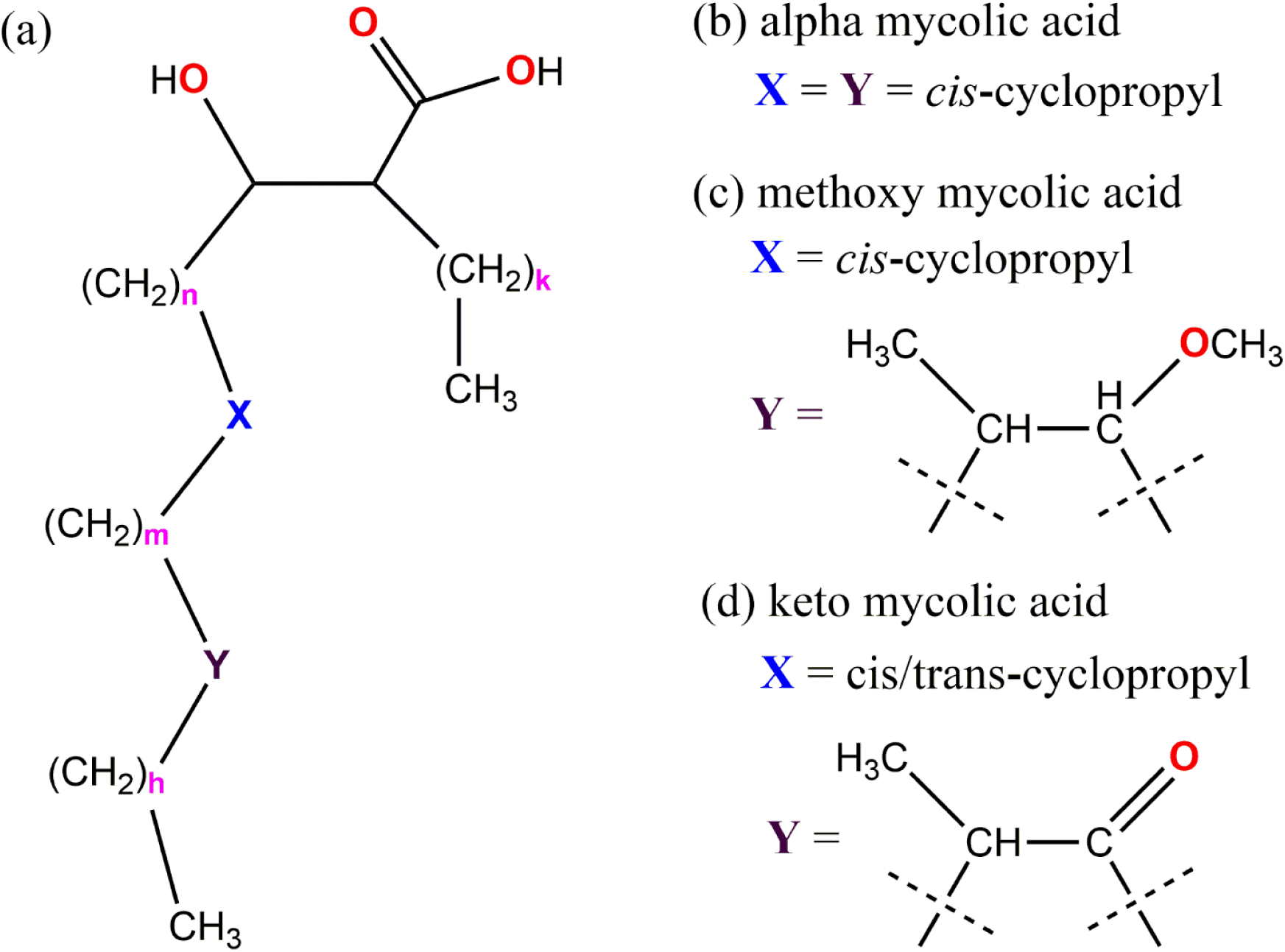
The chemical structure of mycolic acids in *M*. *tb*. cell walls. The typical value of k is 23, commonly denoted as a short chain for all MAs. The extended sequence called the mero chain, composed of the variables n, m, and h, has a cumulative value of 50.^10,21^

The classification of MA is based mainly on functional chain groups, which divide MAs into three main groups: the first group is alpha mycolic acid (AMA), which lacks any oxygen-containing groups within the chain; the second group is methoxy mycolic acid (MMA), characterized by the presence of a methoxy group in the distal position; the third group is keto mycolic acid (KMA), where the distal group of the chain contains a carbonyl group (Figure 1).^10,21^ In *M*. *tb*, the MMA is predominantly associated with *cis*-cyclopropane, while KMA typically exhibits the *trans*-methyl cyclopropane group.^21^ Mycobacterial MAs show notable differences compared to typical FAs. Firstly, they possess a greater length, consisting of 70-90 carbon atoms in total. This length is achieved through the 24 saturated carbons and mero chains^13^ (the elongated molecular segment or long-chain) consisting of 40-60 carbons. Secondly, within the mero chain, functional groups are typically limited to occupying only two specific positions. The proximal position, closer to the β-hydroxy acid, solely carries either cis- or trans-olefin or cyclopropane.

X-ray diffraction experiments show that the MAs form a monolayer in the temperature range of 35° to 41°C, and at higher temperatures, they behave like a liquid.^22^ At temperatures below 35°C, the MAs behave as a plastic solid film, which is very unstable at a lower surface area; however, above this temperature and seven dynes pressure, the monolayer is expanded and more stable.^22^ Hasegawa et al. shown that the AMA forms monolayer at high surface pressure and at sufficient surface pressure the longer alkyl chain of the molecule attains the extended conformation in the monolayer.^23^ In the Langmuir monolayer of KMA, the carbonyl group of meromycolate chain has higher tendency towards aqueous phase leading the molecules to form W-shape conformation.^24^ Moreover, the KMA chain formed a rigid solid condensed monolayer, which was not observed in other MAs chain.^24^ Zhang et al. experimentally demonstrated the triple-folded structure of KMA in the Langmuir-Blodgett monolayer and the carbonyl groups were exposed towards the aqueous phase.^25^ Furthermore, they measured the contribution of hydrophilic and hydrophobic forces over the silicon wafers (coated with hydrophilic film of 1,1,1,3,3,3-hexamethyl-disilazane) surface using colloidal probe force measurement on the MAs monolayer and shown that the electrostatic forces are the main contribution to the long-range repulsive interaction. ^25^ However, the infrared reflection absorption spectroscopy results examines that the meromycolate chain i.e. long chain of the KMA molecules remains folded upon applying the high external pressure to the monolayer surface, while in the AMA monolayer the chains were unfolded.^26,27^ On the contrary, at low surface pressure the meromycolate chain remains in folded conformation for all the MAs in the monolayer.^27^ In the KMA rich monolayer, the mean molecular area was observed much larger than the combined area of AMA and KMA in the mixture, while elastic modulus of the KMA monolayer was close to those of solid.^28^ The mole fraction of AMA trans-cyclopropane that included KMA demonstrated that the α-methyl trans-cyclopropane group stabilized the W-form conformation of MAs in monolayers and solidified them.^28^

The conformational diversity of the long MA molecules arises from the conformational degrees of molecular chains and differences in functional groups. The outer membrane of *M*. *tb*. typically has seven distinct folds of MAs, as illustrated in Scheme 1. MAs conformation is predicated upon the relative disposition of the points a, b, c, d, and e, which are situated at the termini of the chain and correspond to the locations of the functional groups. The conformations above are generally denoted as W, aZ, sZ, eZ, aU, sU, eU and straight.^29,30^ The percentage of these conformations in the monolayer has been studied using molecular dynamics (MD) simulation for various solvent conditions.^30^ However, the temporal evolution in these conformation changes has been reported very recently.^31^ The experimental investigation has demonstrated that these conformations tend to undergo alterations in response to variations in surface pressure, resulting in the acquisition of many folds in the monolayer.^27^ The linear configuration of the AMA within the monolayer was obtained under significant surface pressure, resulting in the monolayer transitioning into a solid state.^23^ Asymmetric distribution of the dipalmitoyl-phosphatidylcholine (DPPC) lipid in the monolayer shows higher stability over a submirosecond simulation time.^32^ However, adding a short cone-shaped tail of diC8PC lipid into the DPPC lipid membrane leads to the release of tension in the asymmetric system by forming a transmembrane pore.^32^ Using the MD simulation, the permeability of the TB-drug molecules was reported through the MA monolayer membrane using the GROMOS and CHARMM force fields and emphasised that the GROMOS FF shows better observation for drug permeability.^33^

Mycolic acids play a pivotal role in eliciting the host’s immune response against the pathogen and have demonstrated efficacy as antigens for the serodiagnosis of tuberculosis.^34,35^ Nevertheless, there still needs to be more understanding regarding the specific organisation of MAs within the cellular wall and their interactions with immunological components. According to experimental findings, the outer bilayer of the mycobacterial cell wall exhibits a width ranging from 7 to 8 nm and displays a symmetrical bilayer configuration.^36–38^ However, the X-ray diffraction studies show that the cell wall lipids are primarily oriented perpendicularly to the surface of the cell wall, potentially resulting in an asymmetrical bilayer structure.^39^ This observation suggests that longer

MAs must be folded to fit this limited space. It has been proposed that MAs exhibit a folding pattern at their functional groups X and Y, as shown in Figure 1, resulting in a W-shaped conformation. Moreover, these MAs are believed to be interposed with lipids in the opposite leaflet by a mechanism resembling a zipper model.^36^ In the groups of oxygenated MAs, it was shown that those with a higher proportion of *trans*-cyclopropane content exhibited a greater propensity to adopt W-form conformations. This suggests that alpha-methyl trans-cyclopropane groups in the meromycolate chain assisted the folding of MAs more than cis-cyclopropane groups.^40^ It is probable that the engagement between the host immune system components, such as antibody binding, entails the presence of a macro-structure comprising several MAs. Hence, understanding the optimal conformations of individual MAs will serve as fundamental units for constructing larger structures and illuminating these domains.

**Scheme 1.**
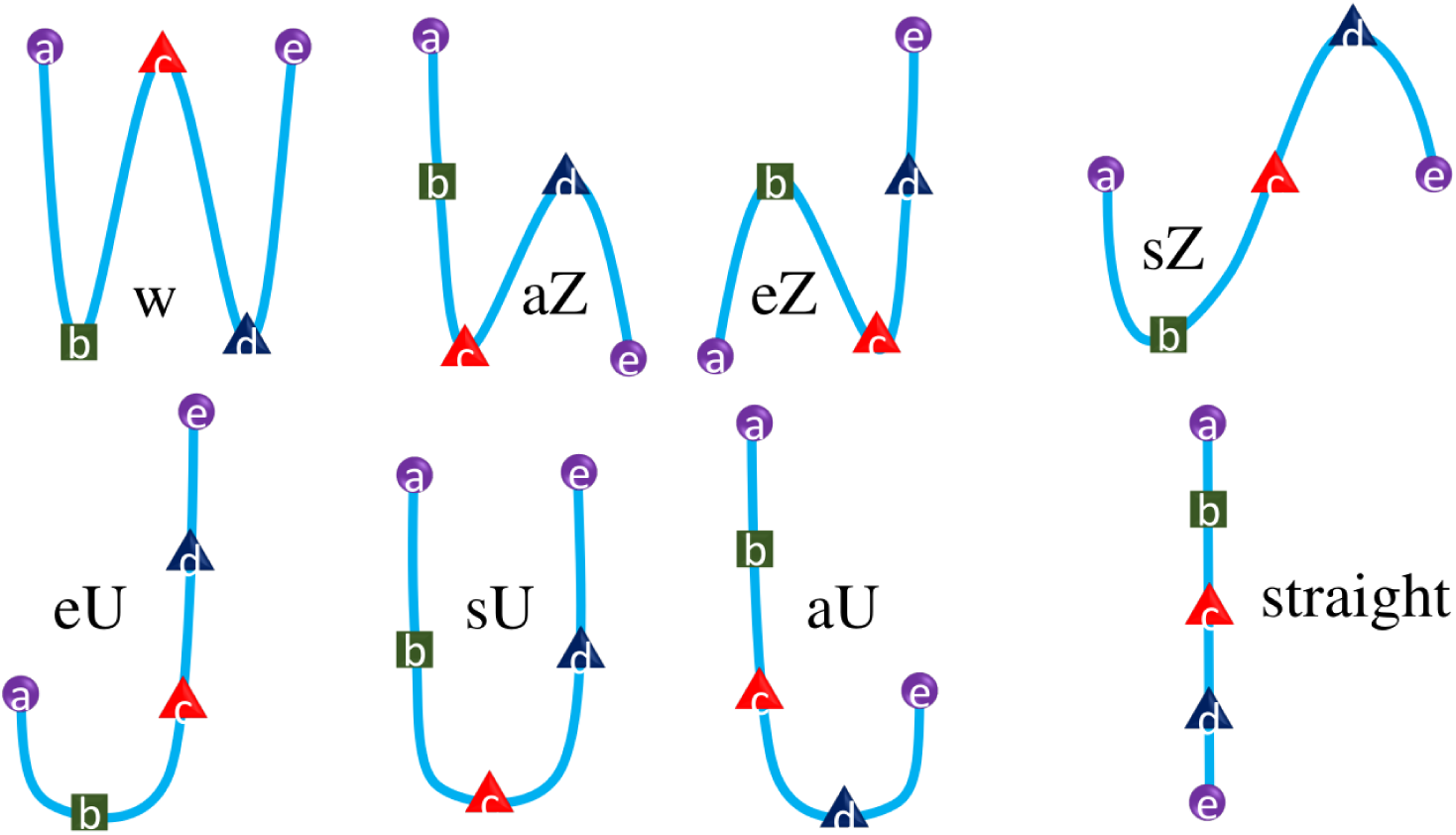
Schematic of seven possible WUZ conformations of mycolic acid in the cell outer membrane.

Experimental evidence demonstrates that the isotherms depicting the relationship between surface pressure and area for AMA, MMA, and KMA exhibit a notable dependence on temperature.^24^ Furthermore, it is seen that the phase diagram of these MAs differs from one another.^24^ The experimental findings also examined that when the temperature and surface pressure are reduced, both KMA and MMA demonstrate the development of rigid condensed monolayers or folded conformations. X-ray diffraction (XRD) results indicate that the MAs exhibit a monolayer structure.^22^ Conversely, at higher temperatures and pressures, AMA, MMA and deoxo-MA tend to adopt elongated conformations. At the same time, KMA is retained in the initial structure.^41,42^ The alpha-methyl trans-cyclopropane unit was crucial for effectively folding the oxygenated MAs.^40^ The conformation of the AMA chain can readily altered, resulting in modification of their W-form in response to the variations in the surface pressure and temperature.^42^

Although the properties of the MA monolayer have been explored experimentally, the insight into the layer formation mechanism and the conformations of MA chains within the assembled layer at a molecular scale is still poorly understood. As described in the previous paragraphs, conformations of MA chains in a monolayer /in solutions have been investigated computationally and all possible conformations have been classified into seven major categories. Despite that, there exist reports on other possible conformations of MA. Moreover, none of these studies addresses the formation mechanism of MA assembly. In the real bacterial cell wall, probable presence of free MA chains in the upper leaflet of mycomembrane together with functionalized Mycolates like TDM/TMM has also been speculated.^10^ Against this background, the major aim of the current study is to observe assembly property of MA using *in silico* tools like molecular dynamics simulation and to probe into the nature of the MA assembly (whether it spontaneously forms a tightly packed monolayer or bilayer) together with a quantitative estimation of MA conformations and the thermodynamical quantities related to assembly formation. This will help to better comprehend the MA layer formation within the *M.tb* bacterium. On the basis of previous experimental observation, there is a lack of understanding in the MAs monolayer membrane formation in the outer cell of *M*. *tb*. Here, we focus to inspect the self-assembly of MAs in aqueous solution. We investigate these membranes in separate single-component and mixture compositions (i.e. AMA = 56%, MMA = 40%, and KMA = 14) that closely resemble the experimentally observed composition of the cell wall. Furthermore, we examine the different MAs’ conformational, structural, and thermodynamic properties separately. The primary objective of this study is to determine whether MAs self-assemble in a monolayer form or bilayer and the conformations and packing arrangements of MAs within self-assembled structure.

## 2. System Setup and Computational Details

The Molecular chains of MAs were built using Avogadro^43^, and the structures were optimised using a generalised amber force field (GAFF). Furthermore, the optimised structures were submitted to an automated topology builder (ATB)^44^ to generate the gromacs^45^ compatible coordinate and associated files. We use the predominant variants of mycolic acids (MAs), specifically alpha-mycolic acid (AMA), methoxy-mycolic acid (MMA), and keto-mycolic (KMA), which were experimentally^4,8,9,19,22^ demonstrated their presence in the *M*. *tb*. outer cell membrane. The elongated MAs were randomly placed in the 9 nm × 9 nm × 18 nm periodic box. Subsequently, the box was solvated using 11000 SPC/E^46,47^ water molecules (100 per MAs). The initial structure of the solvated system is shown in Figure 2.

**Figure 2.**
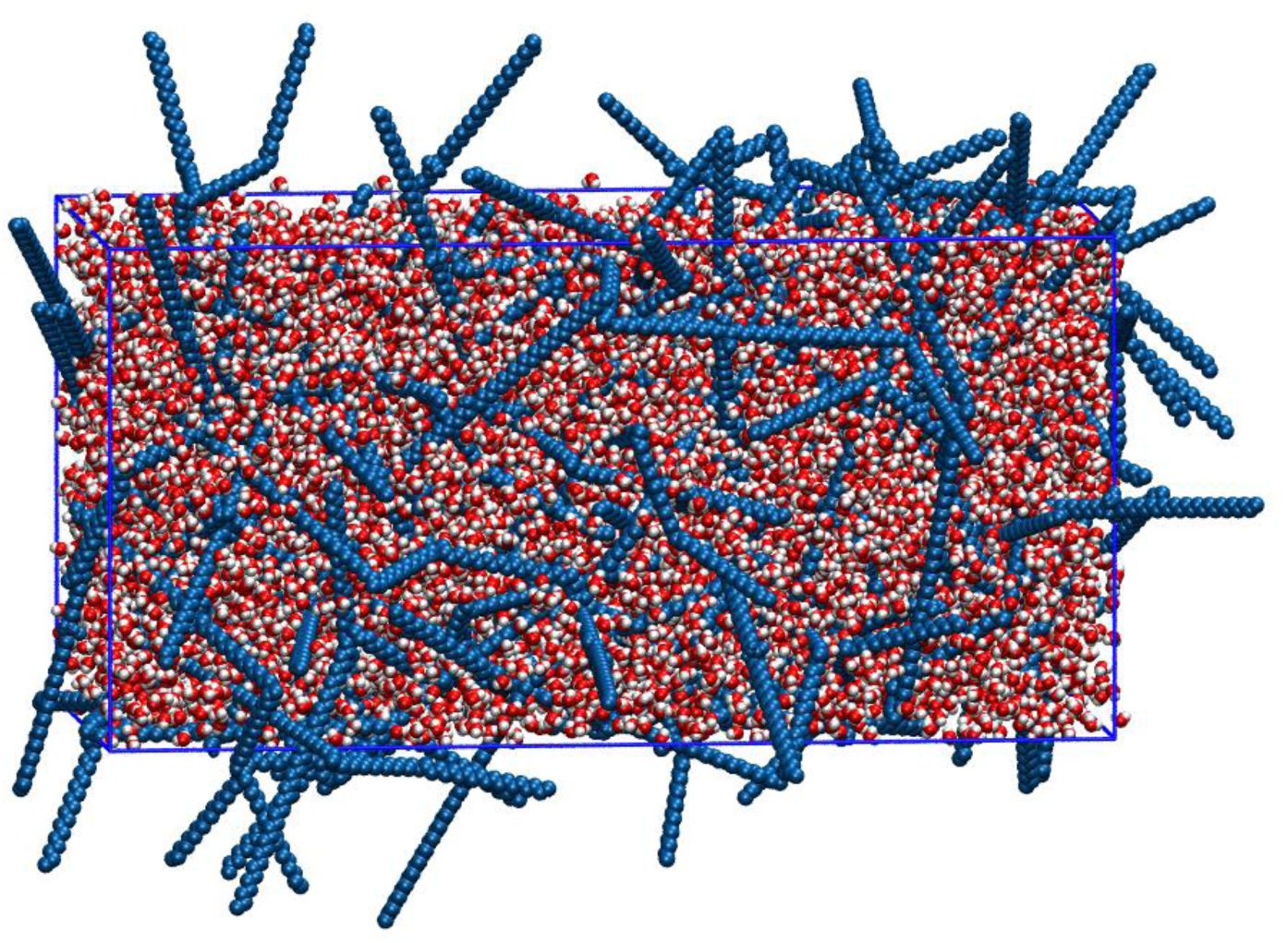
The initial AMA solvated system setup. Colour code: MAs are in blue; water molecules are in vdW representation, with oxygen in red and hydrogen in white.

All the molecular dynamics (MD) simulations are performed in GROMACS^45,48^ (GROningen MAchine for Chemical Simulations) of version 2022.3. Although other force fields are accessible, including CHARMM^49^, OPLS^50^, and AMBER^51,52^, we utilised the GROMOS54a7 force field to describe the inter- and intra-molecular interaction in the mycolic acid.^53^ The force field parameters were obtained from ATB^44^, which provides the DFT-optimized structure and quantum-calculated charges of the atoms in MAs. ^46,47^ The system’s energy was minimized using the steepest-descent method with the convergence criteria of the system energy should be less than 500 kJ/mol. The non-bonded dispersion interactions were computed using the Lennard-Jones 6-12 potential, while the electrostatic interactions were evaluated using the Coulomb potential. The Verlet cutoff scheme was implemented using a grid-based search of non-bonded atoms with a cutoff distance of 1.0 nm. The particle mesh Ewald (PME)^54,55^ method was employed to estimate non-bonded coulomb interaction. All the bonded terms were constrained using the LINCS algorithm^56^ during the MD simulations. All the simulations were carried out at NPT ensemble. The system temperature was maintained 300K using the velocity rescale thermostat^57^ using a temperature coupling constant of 0.1 ps. To obtain the correct density and pressure (1 bar) of the system, we employed the Parrinello-Rahman^58^ barostat isotropically with a pressure coupling constant of 2 ps. The equation of motions was integrated using the leapfrog algorithm with 2 fs timestep. The simulation trajectory was saved every 5000 steps.

## 3. Results and Discussion

### 3.1. Mycolic Acid Monolayer Density Profile

The experimental findings^10,19,22–24,27^ suggest that the mycolic acids in the outer membrane of *M*. *tb*. were arranged in a monolayer form. Few other studies contradict the observation of MAs being present in a monolayer form within the outer cell.^31^ Using molecular dynamics (MD) simulations, we can computationally validate the experimental observations about the development of MA monolayers. From our simulation, it has been observed that the AMA, MMA, and KMA of M. *tb*. MAs in the aqueous solution are assembled as monolayer under isotropic pressure coupling, exhibiting a thickness of ∼ 7 nm, which agrees with the experimental observation.^36^ The MA system self-assembled in a monolayer within 7 to 10 ns of the MD simulation. The structure of the self-assembled monolayer is shown in Figure 3.

**Figure 3.**
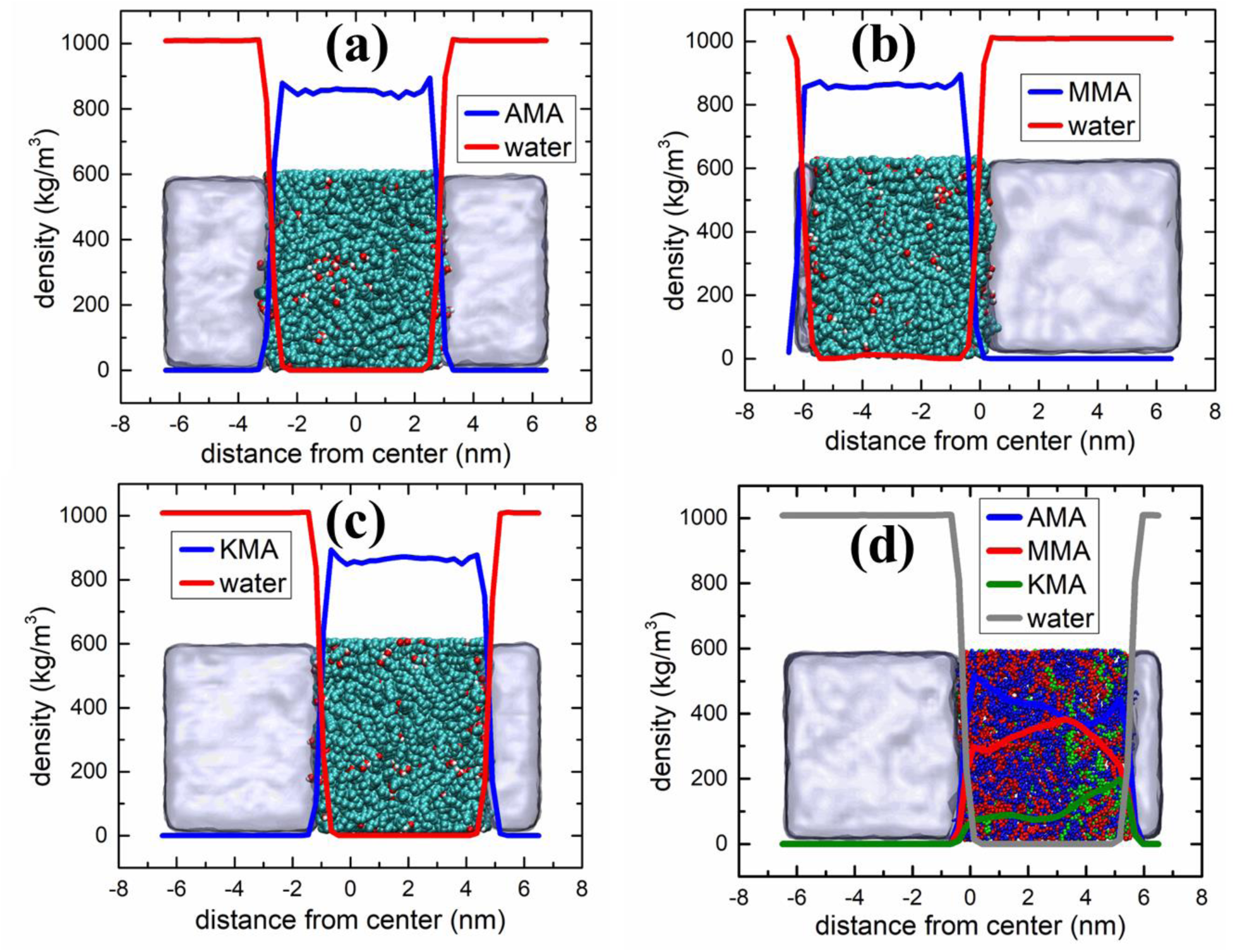
The density profiles (a) AMA, (b) MMA, (c) KMA, and (d) a mixture of MAs (56% AMA, 40% MMA, and 14% KMA) are presented alongside the corresponding water density in each plot. Colour code: water in light ice blue, MAs with vdW representation.

In Figure 3 (a) – (c), we have shown the self-assembled monolayer for all the three different systems (AMA, MMA and KMA). have demonstrated that upon the formation of the MAs monolayer, water molecules exhibit a density of 1000 kg/m^3^ outside the monolayer, while for the single component MAs, the density is 820 kg/m^3^, as evidenced by the enclosed snapshots in Figure 3(a-c). Our simulations for the mixed system consisting of 56% AMA, 40% MMA, and 14% KMA gives the following density values in the self-assembled monolayer form: 420 kg/m^3^ for AMA, 400 kg/m^3^ for MMA, and about 200 kg/m^3^ for KMA. It is evident from Figure 3 that MAs exhibit a monolayer configuration in the outer cell membrane of *Mycobacterium tuberculosis*.

### 3.2. Order Parameters

The internal structure of the MAs in the monolayer (Figure 3) can be characterized by calculating lipid order parameters. We have calculated the order parameter as defined in equ. 1 for the single component and mixed MAs using the gromacs utility (*gmx order*).

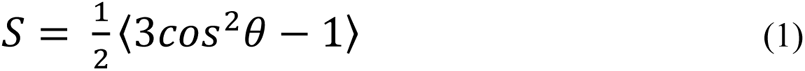

where θ is the angle between the C-H bond vector and the monolayer normal. The angular brackets represent molecular and temporal ensemble averages. The order parameters for the single-component MAs monolayer are shown in Figure 4. The presence of the cyclopropane group in the MAs molecular chain (see Figure 1) causes a deviation in structural order parameters between the 15-20 and 30-35 carbon atoms of AMA and 15-20 carbon atoms of MMA. A sudden drop in order parameter, i.e., S ≈ 0 for the cyclopropane group, suggests its random orientation in the monolayer. In AMA, the short chain is aligned parallel to the monolayer normal, while for MMA, it has a lesser orientation than the AMA. On the contrary, for KMA, the short chain appears randomly oriented in the monolayer. The mero-chain, i.e., long-chain, more ordered structures in AMA and MMA; however, in KMA, monotonic orientations can be seen where S varies from –0.02 to 0.02.

**Figure 4.**
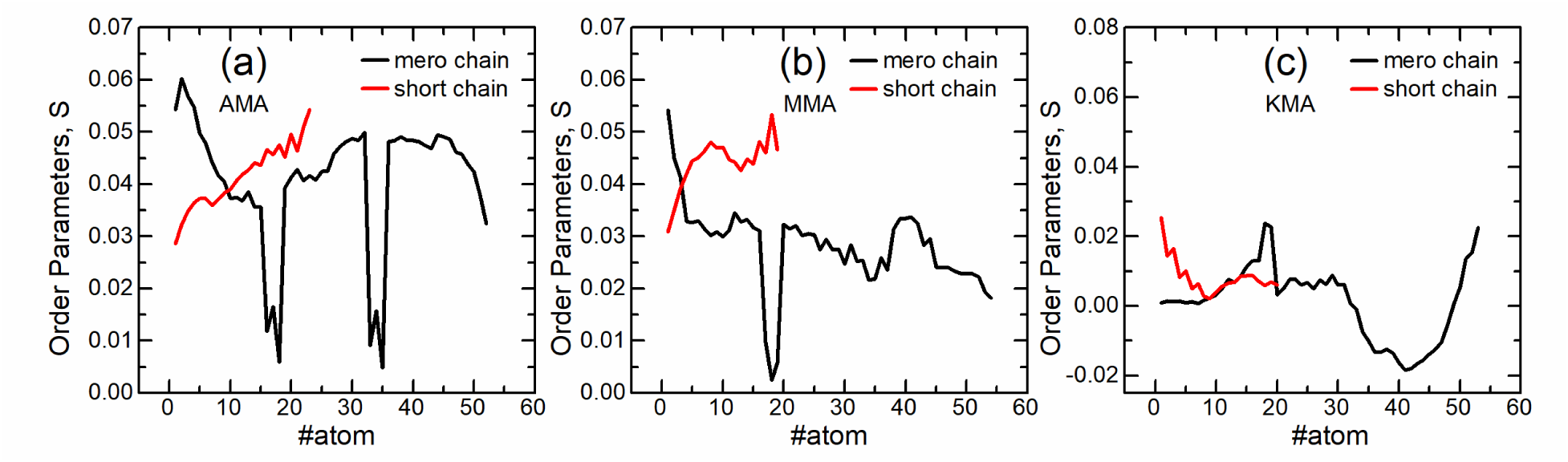
Order Parameter for the MAs (a) AMA, (b) MMA, and (c) KMA in the monolayer.

A random orientation of MAs chains near the cyclopropane group leads the MAs chain to fold in such a manner as to attain a monolayer in the outer cell membrane of *M*. *tb*. Furthermore, we emphasize that the random orientation of such a mero-chain causes the MAs to have different conformations. The percentage of several conformations exits in the monolayer is discussed in the Conformational Analysis Section.

### 3.3. Area-per-Mycolic Acid

The MAs in the monolayer exhibit many conformations and each conformation has a different molecular area. We have calculated the area-per-MAs for the single component MA by averaging the area of all MA divided by the total cross-section of the monolayer over each trajectory frames. The temporal change in the average area-per-MAs of the AMA, MMA, and KMA is plotted in Figure 5. At the beginning of the simulations, the MAs exist in the extended conformations (see Figure 2), leading to more area-per-MAs, as seen in Figure 5 (the enlarged area of the central figure). However, during the MD simulations within five ns of the trajectory, all the MAs self-assembled in monolayer and attain a minimum area/conformation and no change after that throughout the 200 ns to 500 ns trajectory. The MAs achieve the condensed phase upon formation of the monolayer, leading to the constancy in the area-per-MAs over the trajectory. The α-MA exhibits greater area-per-MAs (0.508 nm^2^) as compared to methoxy-MA (0.485 nm^2^) and keto-MA (0.495 nm^2^). Despite the smaller number of atoms per chain in AMA (86) than MMA (91) and KMA (89), the AMA in the MAs monolayer exhibited more are-per-MAs, suggesting lesser packing of it in the monolayer, which is also evidenced from conformational analysis results that the presence of several folds in the monolayer leads to attain a greater area-per-MA. On the contrary, methoxy-MA (MMA) has more excellent packing in the monolayer (Figure 4), leading to the least area-per-MAs compared to other MAs. Furthermore, keto-MA (KMA) exhibits intermediate area-per-MAs in the monolayer.

**Figure 5.**
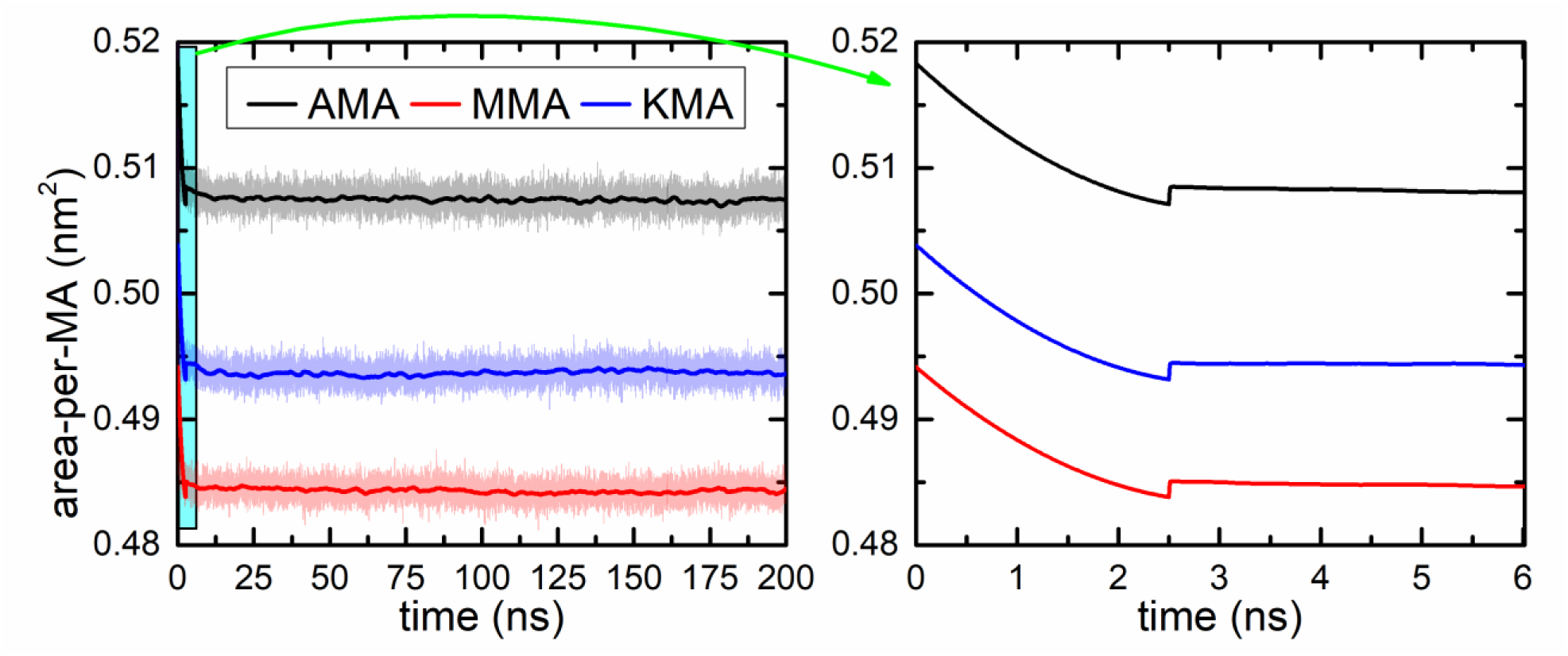
Temporal variation in the area-per-MAs of the α-MA (AMA), methoxy-MA (MMA), and keto-MA (KMA) in the monolayer. (b) Area per MA decreases as a function of time for the initial few ns. We show the initial compaction of the monolayer for the portion (rectangular cyan box) in the first plot is enhanced and presented in the second plot. The plots’ original data (transparent) is smoothed for better clarity in the variations.

### 3.4. Radius-of-Gyration

The structural compactness of the MAs has been estimated by calculating the radius-of-gyration (*R*_g_) for different parts of the mycolic acid chain using the following mathematical expression.

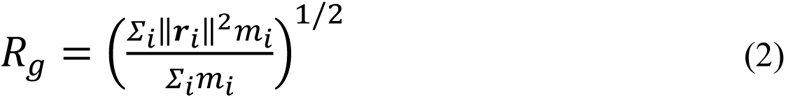

where *m_i_* is the atom’s mass and *r*_i_ is the particle’s position concerning the molecule’s centre of mass. We have divided the mycolic acid chain into three sections ‘*abc’*, ‘*bcd*’, and ‘*cde*’ (see Figure 6A), for better estimation of compactness in the monolayer and calculated the radius-of-gyration (*R*_g_) of each part separately. The results are Molecular Dynamics of Mycolic Acid Monolayers plotted in Figure 6.

**Figure 6.**
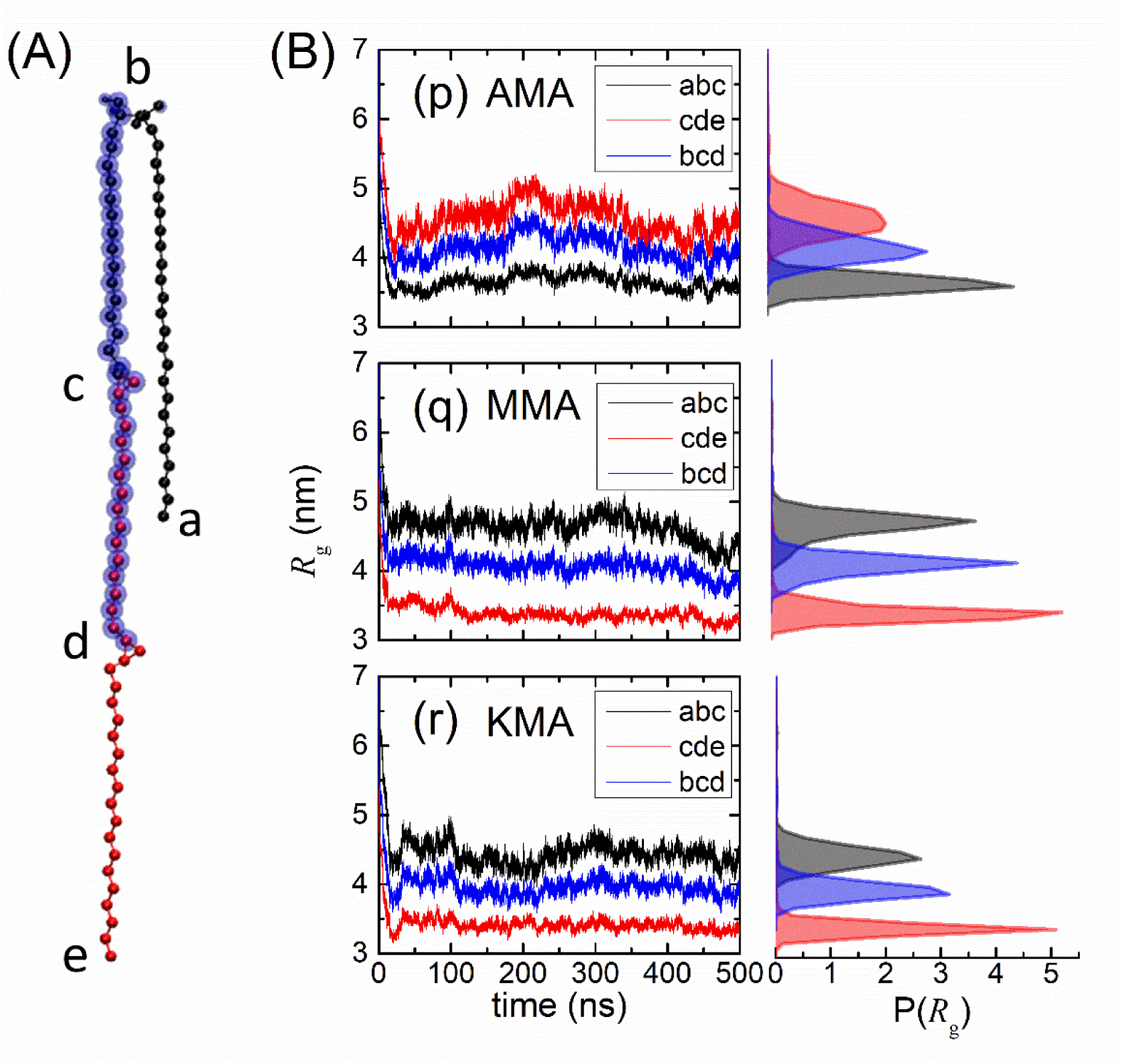
(A) Molecular schematic representation of α-MA chain and its parts. (B) Temporal variation of radius-of-gyration (*R*_g_) of the parts (*abc*, *bcd*, and *cde*) in MAs chains for (p) AMA, (q) MMA, and (r) KMA. Colour codes in (A) are red: section ‘*abc*’, black: section ‘*cde*’, transparent blue: section ‘*bcd*’.

We observed sharp decrease in *R*_g_ for each part of MAs for the initial 20 ns of the simulation trajectory, suggesting the initial adjustment of the chains in the self-assembled monolayer. During the first 20 ns of the trajectory, the MAs chains attain minimum energy conformation; lately, negligible changes have been observed. The size of the *abc* section in α-MA (Figure 6B-p) is smaller than *bcd*, *cde,* designating the short chain in folded conformation (less *R*_g_). Moreover, the mero-chain exhibits higher *R*_g_, causing the AMA chain to have eU and aU conformation at its more probable state, supplementing the conformational probability results discussed in the Conformation Analysis section. For the AMA, apart from seven known conformations such as w, aZ, eZ, sZ, eU, sU, and aU, the population of other conformation (i.e. other than seven) in the monolayer is 72.2%, confirming the random and extended conformation of the chains in the assembly which further suggest the lesser packing of the chains in the monolayer. At the same time, for MMA and KMA monolayers, it is 69.9% and 64.68%, respectively, which is lesser than AMA due to the reduced size of the *cde* part of the chains. A reversal in *R*_g_ can be seen for *abc* and *cde* parts of the methoxy- and keto-MA in the MAs monolayer (Figure 6B-q & 6B-r). AMA’s ‘cde’ parts could not pack densely because of two cyclopropane groups in its mero chain at position ‘c’ and ‘d’ (see Figure 1 and Figure 6A). In contrast, the absence of one cyclopropane group (see Figure 1) in mero chains of KMA and MMA enables the ‘*cde*’ part of these two molecules to fold more compactly (Figure 6B-q & 6B-r).

The probability distribution of *R*_g_, P(*R*_g_) for all the *abc*, *cde*, and *bcd* parts of the MAs ar e shown right to the Figure 6B. The *cde* and *bcd* parts of the AMA chains are in broad distribution as compared to *abc*, suggesting the flexible nature of the chains at their functional group (i.e. cyclopropane group) (see Figure 1). The *abc* part in the AMA shows rigid behavior as can be seen from the temporal change in *R*_g_ and sharp distribution of the P(*R*_g_). However, for the case of MMA and KMA, a rigid nature also observed in all the parts of the MAs chains. Therefore, it can be deduced that the AMA has more significant flexibility in the assembly than that of the MMA and KMA mycolic acid chains.

### 3.5. Conformations Analysis

Mycolic acids (MAs) in the *M*. *tb*. outer-cell membrane exhibit seven different WUZ conformations, namely w, aZ, eZ, sZ, eU, sU, and aU, as revealed from several combined studies of MD simulations^11,30,41^ and experiments.^22,23,27,42^. We performed atomistic MD simulations of 110 MAs chains in an aqueous environment and calculated the percentage of WUZ conformation for single component (i.e. AMA, MMA, and KMA) and mixture cases (i.e. 56% AMA, 40% MMA, and 14% KMA). Each frame of the simulated trajectory was analysed for the seven possible WUZ conformations based on their intermolecular distances among the several parts of the MAs (see Figure 6A). The intermolecular distances between these parts (a-c, a-e, c-e, and b-d) were taken from Groenewald, W. et al.^30^ The conformational snapshots of seven possible WUZ and straight conformations obtained from MD simulations are shown in Figure 7. The percentage fold of each MAs in a single component and mixture was obtained from the self-written Python code. The results are plotted in Figure 8.

**Figure 7.**
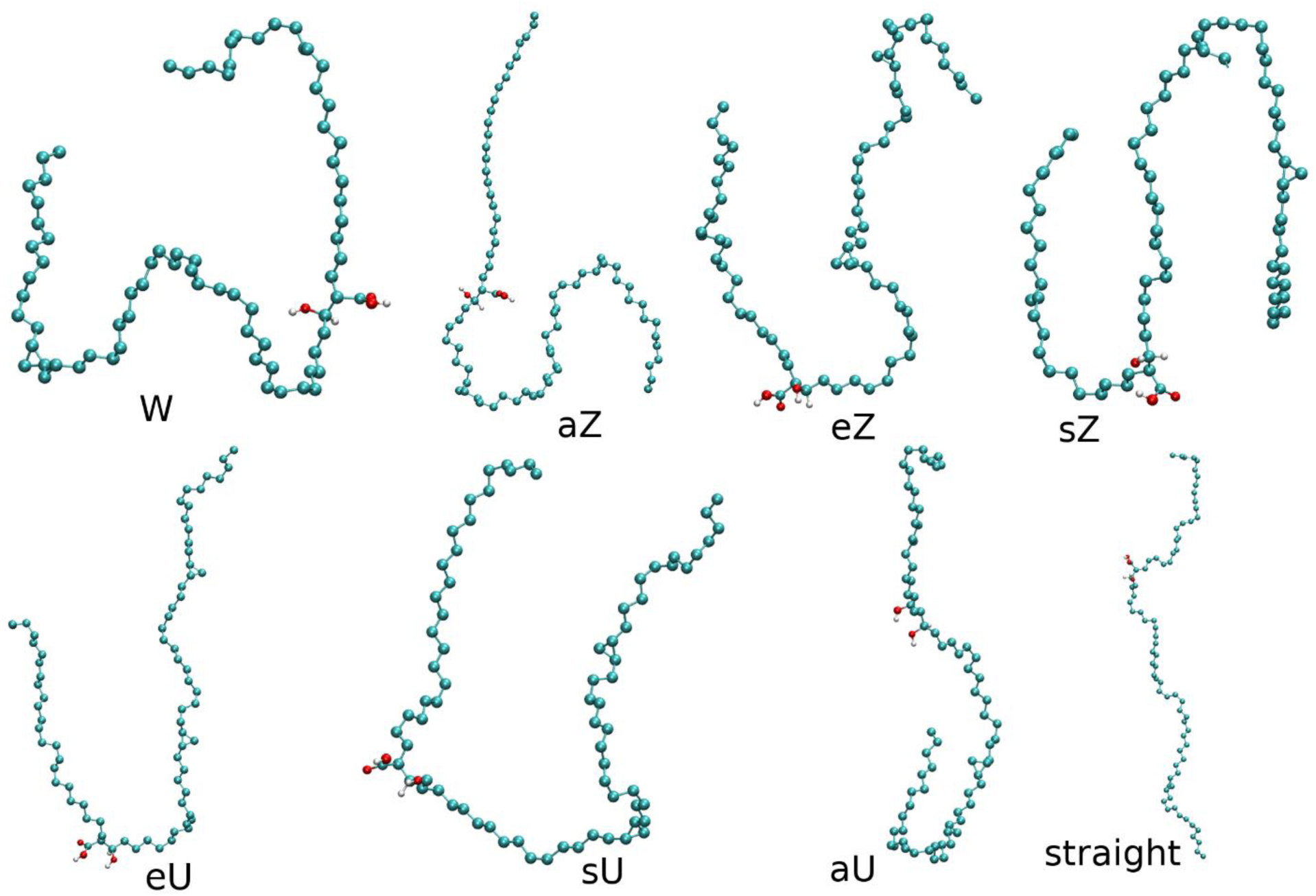
Conformational snapshots of average structures of α-MA (in single component) in its seven different WUZ-folds in the monolayer.

**Figure 8:**
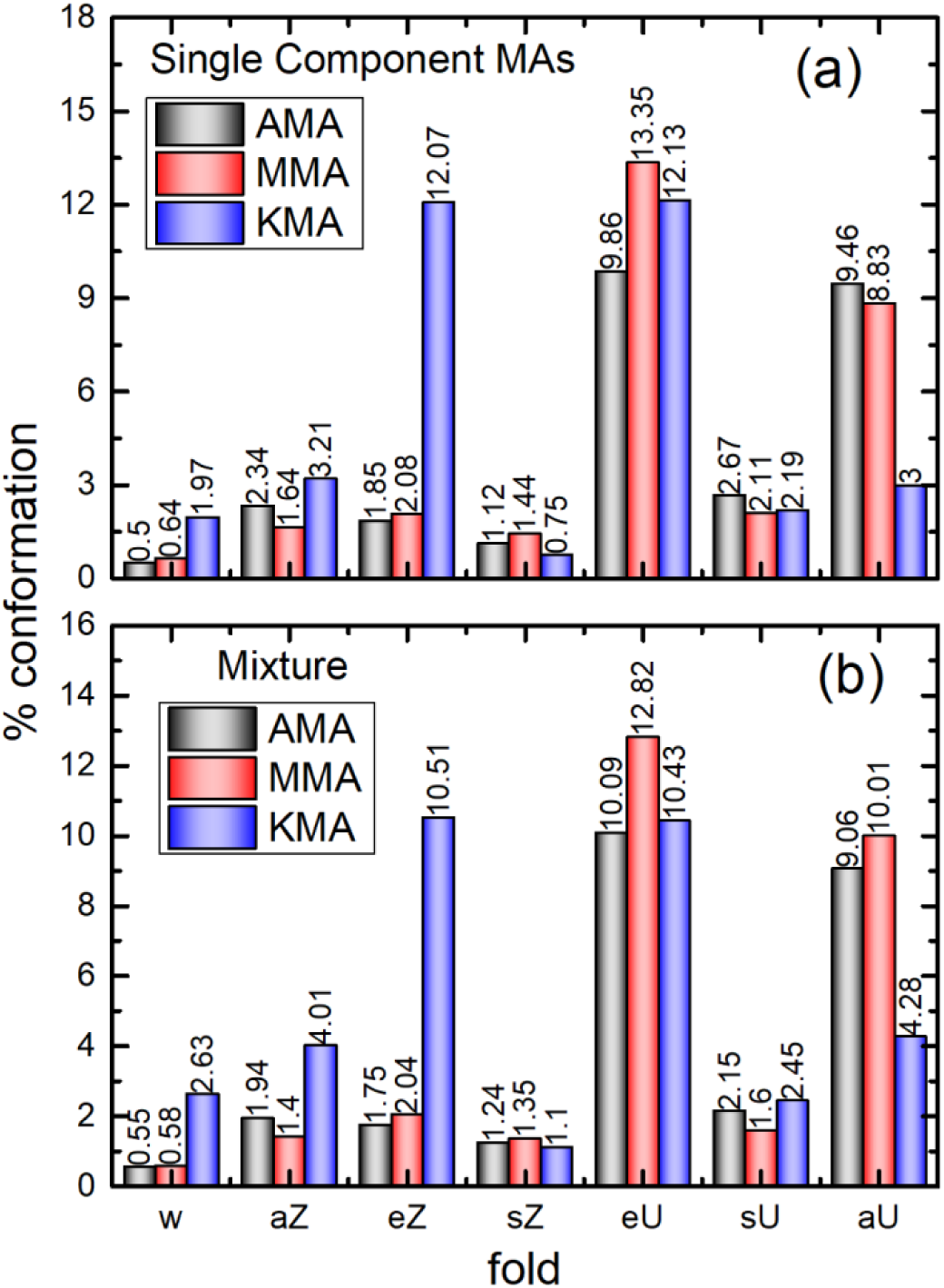
Percentage of conformation of several folds of MAs in (a) single-component of the AMA, MMA, and KMA, (b) in their mixture with 56% AMA, 40 % MMA, and 14 % KMA composition.

For KMA, the highest percent conformation for eZ is observed, and overall eU folds exhibit the most significant population for all α-MA, MMA, and KMA compared to other folds in the monolayer (Figure 8a). The keto group at the distal position of the KMA chain suppresses different folds except for the eZ because the oxygen molecules prefer to stay in the water-monolayer interface. Therefore, the KMA chain adopts the monolayer’s more stable or favourable folds (eZ and eU). In contrast, the W fold in the single component and mixture retains the lowest percent population (0.68% and 0.54%, respectively), followed by the sZ fold (1.44% and 1.35%, respectively), stays the lowest population in the KMA monolayer. In methoxy-MA, the eU fold exhibits the highest percent population (13.35% in single component and 12.82% in combination), followed by aU (8.83% in single component and 10.01% in combination) fold in the MMA monolayer.

The aZ fold also exhibits more percent conformation in the KMA (3.21% in single component and 4.01% in mixture) compared to α-MA (2.34% in single component and 1.94% in combination) and MMA (1,64% in single component and 1.40% in combination). Therefore, it is clear that the KMA in both single component and mixture exhibits a higher probability of attaining the extended folds (namely aZ, eZ, eU, and aU) of the seven different conformations in the monolayer. On the contrary, in α-MA and MMA, despite a higher probability in eU and aU, the other conformation (folds) are also present to make the monolayer more impermeable to the drugs.

The conformations (folds) that are more folded (coiled), such as W, sZ, and sU, may cause the MAs to be more impermeable than extended folds. The distribution of intermolecular distance of parts (see Figure 6A) of MAs for the other conformation is plotted in Figure 9. The intermolecular distance of the parts for the other conformation is higher than that given in Ref. 22 and exhibits a higher percentage in both the single component and mixture monolayer. It is evident from Figure 8 that the seven conformations in single component for the α-MA, MMA, and KMA exhibit 27.8%, 30.1%, and 35.32%, respectively, while other conformations exhibit 72.2%, 69.9%, and 64.68%, respectively. Therefore, the other conformations are more likely in the MAs monolayer. However, in a mixture with 56% AMA, 40 % MMA, and 14 % KMA composition, α-MA (or AMA), MMA, and KMA exhibit 26.78%, 28.40%, and 35.41%, respectively and the other conformation stays 73.22%, 71.6%, and 64.59%, respectively. The distribution of the intermolecular distance between the ‘a’ and ‘e’ (Figure 6A) given in Figure 9 (I & II) suggests that for the other folds, this segment has inclusive values ranging from 0.25 nm to 8 nm over the trajectory, while for the ac, ce and bd, most of the intermolecular distances range from 2 nm to 6 nm which are unable to come in the range WUZ conformation definitions.^30^

**Figure 9:**
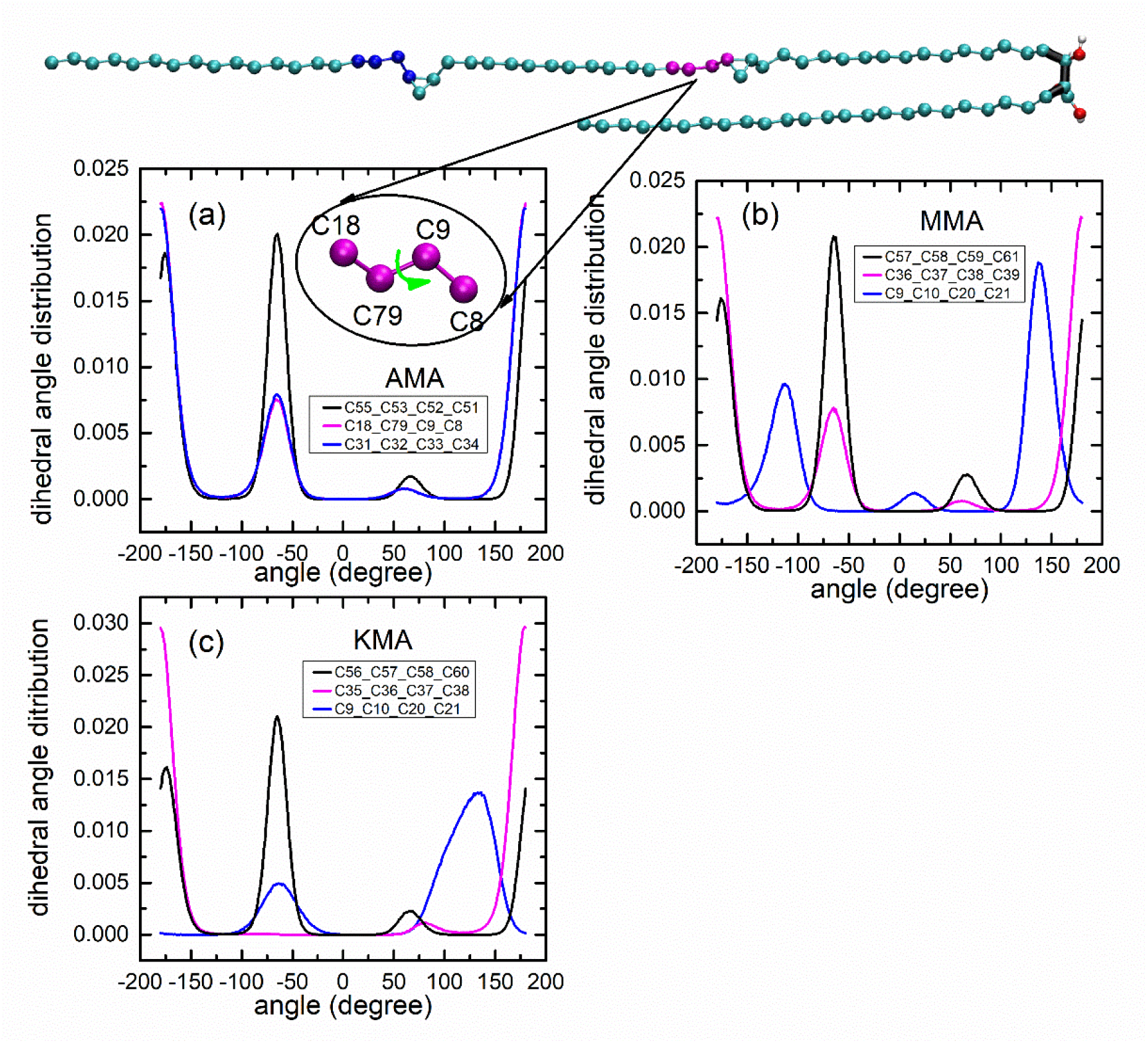
The average dihedral angle of the mycolic acids adjacent to cyclopropane and at the functional group for (a) AMA, (b) MMA, and (c) KMA in single component monolayer. The torsion angle adjacent to the cyclopropane group is given in magenta and blue, while blue is for the torsion angle between the -COOH and -OH groups of the MAs chains.

The other fold in Figure 10 of the MAs monolayer is impotent to attain the WUZ conformation in 200 ns to 500 ns of the MD simulation trajectory. Some MAs chains for the *ac*, *ce*, and *bd* segments (Figure 6A) are in 4 nm to 7 nm intermolecular distance, which suggests that several other folds are in straight (or extended) conformation, and this range is more probable for the keto-MA (Figures 10 C&F). It is also essential to iterate here that the WUZ convention used to describe the conformations of mycolic acid chains only considers the intramolecular distances, not the change in the dihedral angles.^31^ Besides the variation of the intramolecular distances beyond the WUZ convention, the dihedrals of the MA chains also altered significantly during the course of our simulation, and as a result, most of the mycolic acid chains folded beyond the usual WUZ conformational shapes. Hence, an abundance of other conformations prevailed.

**Figure 10.**
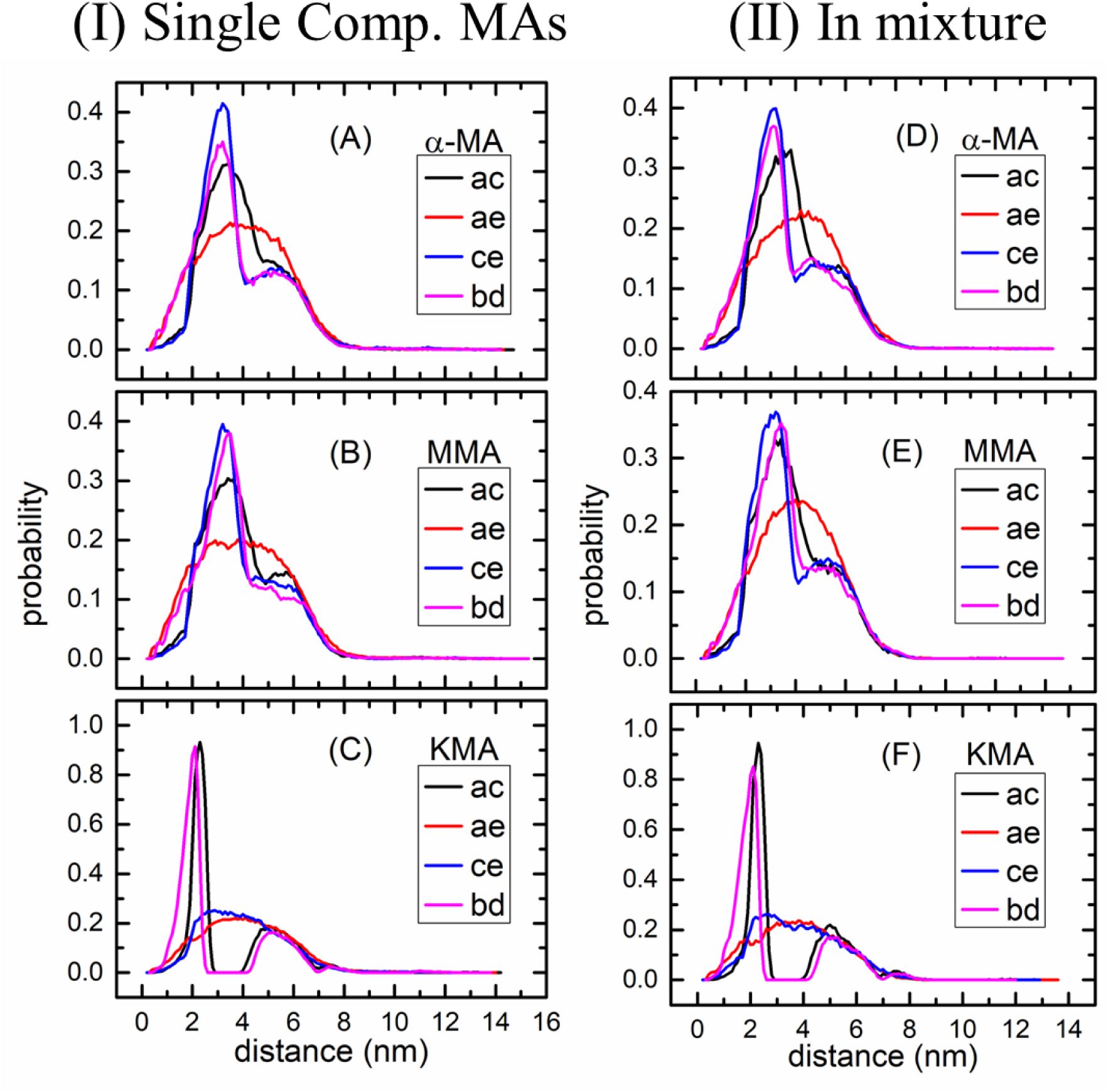
The distance distribution of ac, ae, ce, and bd calculated from the equilibrium sampling period of the trajectory for (I) the single component MAs in the monolayer and (II) distribution in the case of mixture with 56% AMA, 40 % MMA, and 14 % KMA composition.

Furthermore, the torsion angle distribution adjacent to the functional groups is calculated, and the results are plotted in Figure 9. The dihedral angle connecting the carboxylic group (–COOH) and a hydroxyl group (–OH) of the MAs shows a higher probability in *gauche*ˉ (*g*ˉ) (conformation, causing the chains to fold/coiled), which is observed to occur at –60° suggesting the MAs chains to folds very frequently at this group. This observation also supplements the percent conformation (Figure 8) calculated for the MAs chains in the monolayer. The eU folds in the monolayer are in the decreasing order of MMA > KMA > AMA (Figure 8). Moreover, the *trans* (±180°) conformations also show pronounced distribution. For the AMA in Figure 9(a), the *trans* conformations have a greater probability (color: blue and magenta), leading to higher eU and aU folds for the AMA in the monolayer (Figure 8).

The absence of a cyclopropane group (see Figure 1) in the MMA and KMA shows a different dihedral angle distribution than the AMA. The dihedral distribution near the proximal position’s cyclopropane, i.e., C35-C36-C37-C38 (magenta color) in the KMA, shows the *trans* conformation at the highest probability, suggesting most of the KMA chains in the monolayer exhibit the eU (an extended conformation) folds with 12.13 % in single-component mycolic acid and 10.43% in the mixture (Figure 8). However, in the MMA single-component monolayer, the same dihedral, i.e., C36-C37-C38-C39, shows a *g*ˉ (–60°) peak at the significant intensity due to the cyclopropane group leading to form several folds such as W, aZ, eZ, and sU in the monolayer. Their percentage populations can be seen in Figure 8. Moreover, the functional groups at the distal position for the MMA and KMA are methoxy and keto, respectively, and their dihedral (C9-C10-C20-C21 in blue) distribution peaks appear at ±120° for the MMA and –60°, +120° for the KMA. The orientation and twisting of this dihedral lead to the formation of those folds that bend at the *d* point in the chains (Figure 6A).

Another significant feature of Figure 10 is the slightly narrow distribution of the intramolecular distances for KMA compared to the other two types of MA chains (Figure 10 (c) and 10(F)). The probable reason behind this characteristic is the presence of the keto group in the distal position of the mero chain of KMA. The oxygen has an affinity toward water, which limits the folding ability of the KMA molecule, giving rise to a narrower intramolecular distance distribution.

### 3.6. Principle Component Analysis

In order to obtain a better insight into the stability of various mycolic acid molecules inside the self-assembled monolayer, we have performed the principal component analysis of the system and computed the free energy landscape w.r.t PC1 and PC2 (Figure 11). From Figures 11 (a)-11(c), it is evident that there exist large valleys with minimum free energy value for AMA and KMA in single component assembly (Figures 11(a) and 11(c)). A slightly smaller valley was observed for MMA (Figure 11(b)). As PCs capture fluctuations of the systems, the presence of a large basin in the free energy landscape indicates that significant structural and spatial fluctuations of the mycolic acid molecules took place inside the stable assembly within the course of the simulation. A notable observation is that, in the case of multi-component assembly, the size of the valley with minimum free value contracted which is more significant for AMA and KMA (Figures 11(d)-11(f)). This indicates a comparatively restricted movements of the mycolic acid molecules inside the stable monolayer assembly when all kinds of mycolic acids are present together.

**Figure 11.**
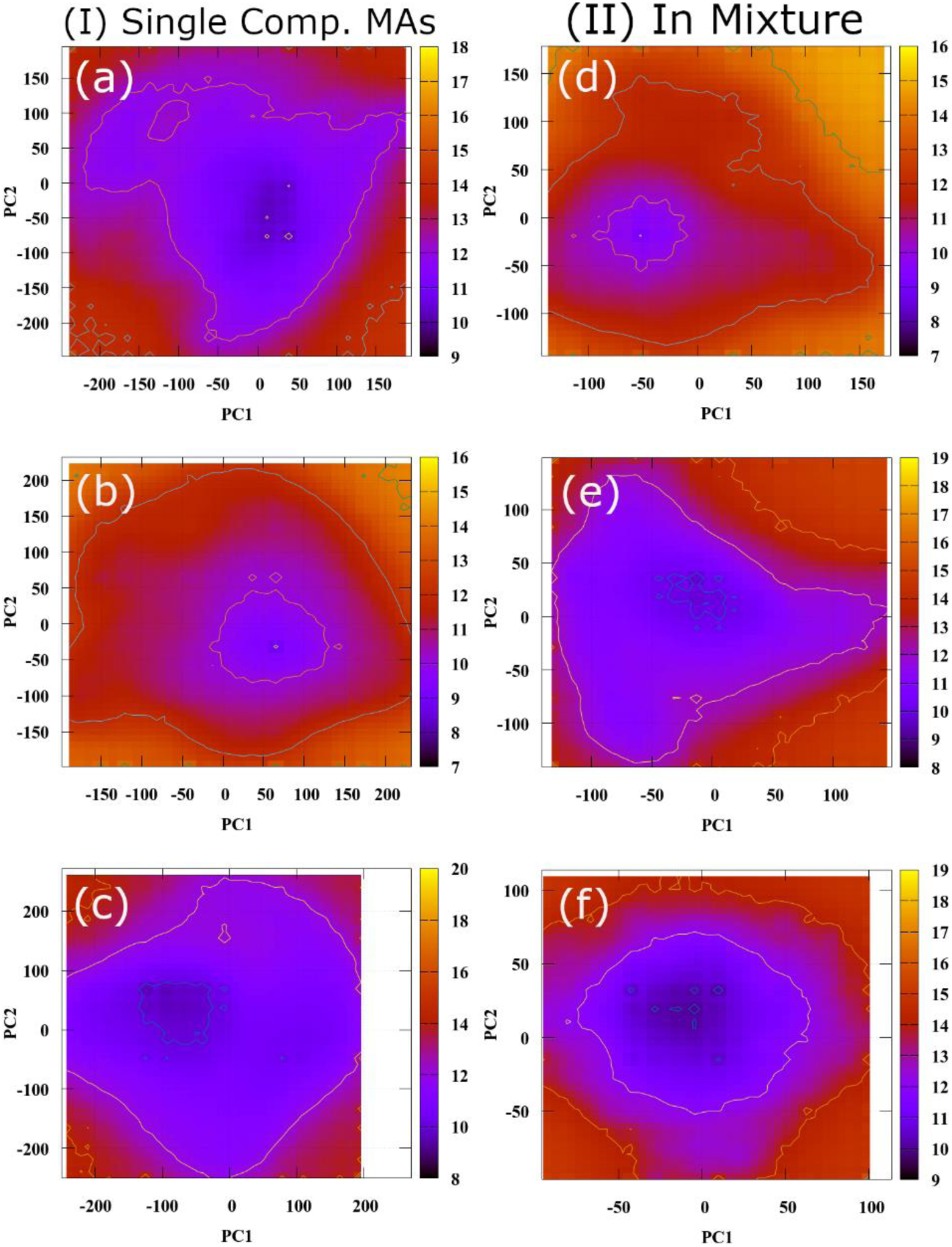
Free-energy-landscape (FEL) for (a) α-MA, (b) methoxy-MA, and (c) keto-MA in the single component monolayer and respective MAs in the mixture is given in (d-f).

We have also calculated the Free energy landscape of the monolayer assembly w.r.t the average RMSD and radius of gyration of the mycolic acids (Figure 12). From Figures 12(a)-12(c), it is evident that in pure monolayer assembly, the free energy minima for AMA is located at high *R*_g_ and low RMSD region, whereas the same for MMA and KMA lies at high RMSD and low *R*_g_ region. These observations indicate that in the pure monolayer assembly, AMA packs less densely within the monolayer than MMA and KMA because the ‘*cde*’ part of AMA folds less effectively than KMA and MMA (Figure 6). Significant alteration of this behaviour was recorded for mixed monolayer (Figures 12(d)-12(f)). In this case, free energy minima were found to be located in a similar region for all three types of MA chains (Figures 12(d)-12(f)). In corroboration with the previously stated observations, this specific change in the free energy landscape of AMA in mixed monolayer implies that the AMA chains become more stable and densely packed in the presence of other MA chains.

**Figure 12.**
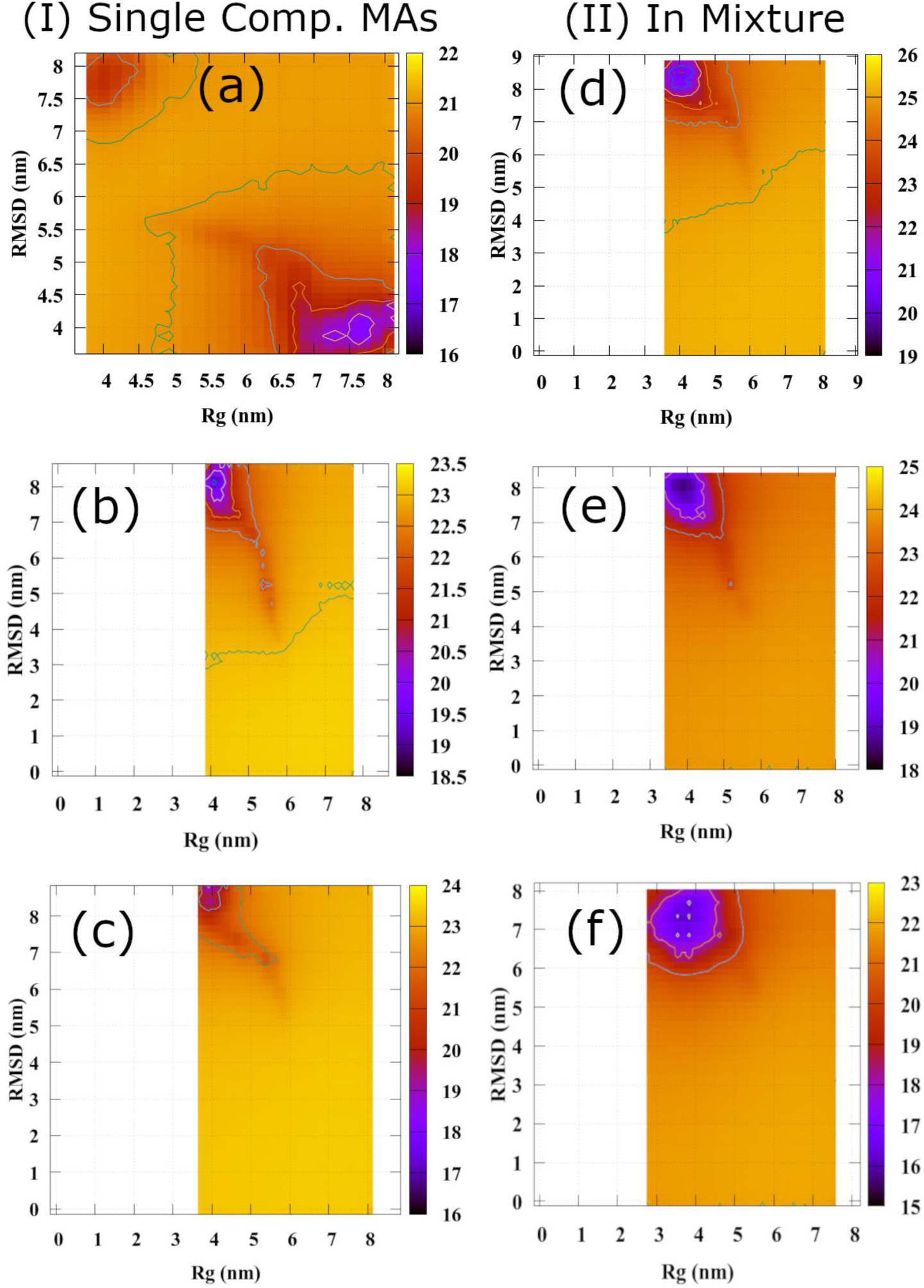
Free energy landscape (FEL) of the mycolic acid monolayers as a function of RMSD and *R*_g_ in (I) single component MAs for (a) AMA, (b) MMA, and (c) KMA and the respective FEL for the MAs in the mixture are from (e-f).

### 3.7. Thermodynamics of monolayer formation

The temporal change in non-bonded energy between the mycolic acids and solvent (i.e., water) molecules has been calculated, and the results are plotted in Figures S2-S4 (Supporting Information). We found that the van der Waals (vdW) energy is more dominating for the assembly formation. Initially, the vdW energy associated with mycolic acids and water molecules is more negative (∼12000 kJ/mol), suggesting a more significant interaction. At the same time, after the monolayer formation (within 20 ns of the trajectory), the MAs chains have more intermolecular interaction with the other MAs chains in the monolayer than their interactions with water molecules. This observation can be seen from the non-bonded energy plots given in Figures S2-S4 for the AMA, MMA, and KMA. However, Coulombic energy among the MAs and water molecules has a less significant role in the assembly formation, which shows negligible changes over the time within 20 ns of the trajectory compared to vdW.

Additionally, MAs’ free energy landscape (1D) was calculated concerning other physical quantities to obtain better clarity in the thermodynamic properties of the single-component monolayer. To achieve this goal, we performed the metadynamics^59^ simulations (600 nm to 800 nm) using Plumed^60^ (an open-source platform) patched with GROMACS.^45^ A distance-based collective variable (CV) is used between the first two chains in the monolayer. The Gaussian width and height are set to 0.005 nm and 0.05 kcal/mol, respectively, for the biasing of the z-component of the centre of mass (com) between two chains. The results obtained are plotted in Figure 13. The keto-MA exhibits the lowest free energy minima (ΔG = -125 kcal/mol) over others, while methoxy-MA shows the highest (ΔG = -92 kcal/mol). On the contrary, in the alpha-MA, the lowest is obtained at ΔG = -102 kcal/mol) over several energy minima in the landscape. One notable characteristic of the free energy landscape is the appearance of several prominent free energy minima, specifically for AMA and MMA (Figure 13), which can be attributed to the broad distribution of ‘*ac*’ and ‘*bd*’ intramolecular distances found for AMA and KMA (Figure 10(I)(a)-10(I)(c)). As stated, a broader distribution of intramolecular distances indicates that more stable configurations can exist in the assembled monolayer, leading to more than one free energy minimum in the energy landscape (Figure 13).

**Figure 13.**
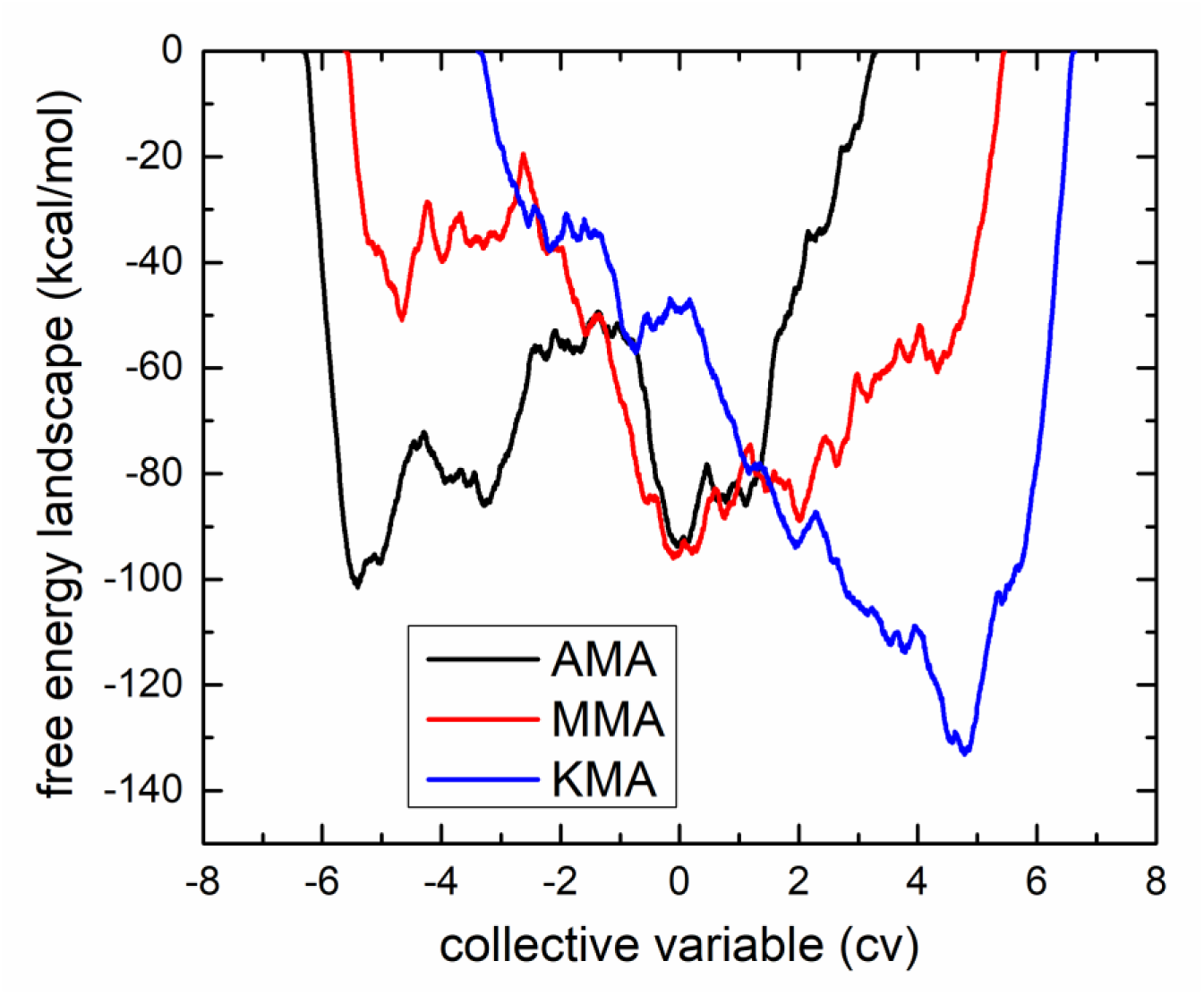
The Free energy landscape of the MAs in a single component monolayer obtained from enhanced metadynamics sampling simulations.

In order to obtain a clear idea about the thermodynamic state of the assembled monolayer system, we have plotted the time evolution of the distance between the two mycolic acids, which has been used as the CV in metadynamics simulation (Figure S1 in Supporting Information). From these figures, we can conclude that thermodynamically, all pure assembled monolayers attained local free energy minima (Figure 13 and Figure S1 in supporting information). Hence, all assembled monolayers are thermodynamically stable. However, we speculate that a long time will be required for these systems to attain global free energy minima.

It is clear from the free energy landscape that the greater the free energy minima, the lesser the chance for any molecules to pass through the monolayer. Therefore, the KMA has more energy barrier for the drug to penetrate through the monolayer.

The AMA exhibits the intermediate resistance (based on free energy value) for the drug to pass through the monolayer. All seven folds in the AMA monolayer (Figure 8) with a higher population of eU and aU conformations show that the free energy has several valleys in the landscape, which is also true for the methoxy-MA (MMA).

### 3.7. Conclusions

We have simulated the *M*. *tb*. outer cell membrane component *Mycolic acid* for better clarity and observance of monolayer formation in aqueous media. Our results demonstrate that functional groups in the mycolic acid enable the folding in the seven different conformations in the monolayer. The MAs attain the condensed phase monolayer of 5 nm to 6 nm thickness within five ns of the trajectory and stay over the course. For the mixture of mycolic acid with 56% α-MA (AMA), 40 % methoxy-MA (MMA), and 14 % keto-MA (KMA) composition, the results show that MMA has more preference to stay in the middle of the monolayer in comparison with AMA and KMA due to presence of more hydrophobic methoxy-group in the distal position of the chain. In a single-component mixture, the monolayer density was 820 kg/m^3^, while based on the available composition of the MAs in the mix, the density obtained was 420 kg/m^3^, 400 kg/m^3^, and approximately 200 kg/m^3^ for the AMA, MMA, and KMA, respectively.

The short chain has a more ordered structure in the monolayer than the mero-chain (long-chain). In contrast, the mero-chain in the AMA and MMA has more deviation in the ordered structure at the cyclopropane functional group. The area-per-MA for the single component was observed to decrease in the order of AMA > KMA > MMA. The least area-per-MA of the MMA suggests the formation of a more compact monolayer with all possible folds. The size of the ‘*abc*’ segment of the AMA is lesser than the MMA and KMA, indicating that the part of the chain has a more compact structure in the monolayer, which has a significant role in forming WUZ conformation. The percentage of WUZ conformation results revealed that all seven possible folds are present in the monolayer for AMA, MMA, and KMA. For KMA, the most favourable folds are eZ and eU, with 12.07 % and 12.13 % of their population. However, eU and aU conformations are more probable in AMA and MMA for both the single component and mixture. The intramolecular distance distribution between the segments of MAs for the other folds (other than seven folds) was calculated and plotted. The average value is observed to be greater than the required for estimation of folds.^30^

Change in free energy with the changing COM-COM distance between two MA chains was computed using metadynamics simulation, which illustrated that upon assembling, all systems reached local minima in the free energy landscape, pointing toward the thermodynamic stability of the assemblies.

In summary, the current study provides molecular-level insight into the experimentally observed assembly property of mycolic acid chains, establishes their thermodynamic stability, and dissects the conformational behavior of MA chains during assembly formation.

## Acknowledgement

The authors thank the Department of Biotechnology (DBT), Government of India, for funding this work (Grant number: BT/PR33123/MED/29/1497/2020). We also recognise the computational resources in our group where all the simulations were performed. The molecular modeling and coding were carried out in the computational facility provided in PKMLab at the Center for condensed matter physics, department of Physics, IISc Bangalore, India.

## Notes

### Competing Interest Statement

The authors have declared no competing interest.

## References

(1) Global Tuberculosis Report 2022; World Health Organization, 2022.

(2) Zeigler, D. R.; Perkins, J. B. The Genus Bacillus. In Practical handbook of microbiology; CRC Press, 2021; pp 249–278.

(3) Jarlier, V.; Nikaido, H. Mycobacterial Cell Wall: Structure and Role in Natural Resistance to Antibiotics. FEMS Microbiol. Lett. 1994, 123 (1–2), 11–18. 10.1111/j.1574-6968.1994.tb07194.x.

(4) Brennan, P. J.; Nikaido, H. The Envelope of Mycobacteria. Annu. Rev. Biochem. 1995, 64, 29–63. 10.1146/annurev.bi.64.070195.000333.

(5) Trias, J.; Benz, R. Permeability of the Cell Wall of Mycobacterium Smegmatis. Mol. Microbiol. 1994, 14 (2), 283–290. 10.1111/j.1365-2958.1994.tb01289.x.

(6) Jarlier, V.; Nikaido, H. Permeability Barrier to Hydrophilic Solutes in Mycobacterium Chelonei. J. Bacteriol. 1990, 172 (3), 1418–1423. 10.1128/jb.172.3.1418-1423.1990.

(7) Liu, J.; Barry, C. E.; Besra, G. S.; Nikaido, H. Mycolic Acid Structure Determines the Fluidity of the Mycobacterial Cell Wall. J. Biol. Chem. 1996, 271 (47), 29545–29551. 10.1074/jbc.271.47.29545.

(8) Dulberger, C. L.; Rubin, E. J.; Boutte, C. C. The Mycobacterial Cell Envelope — a Moving Target. Nat. Rev. Microbiol. 2020, 18 (1), 47–59. 10.1038/s41579-019-0273-7.

(9) Batt, S. M.; Minnikin, D. E.; Besra, G. S. The Thick Waxy Coat of Mycobacteria, a Protective Layer against Antibiotics and the Host’s Immune System. Biochem. J. 2020, 447 (10), 1983–2006. 10.1042/BCJ20200194.

(10) Marrakchi, H.; Lanéelle, M. A.; Daffé, M. Mycolic Acids: Structures, Biosynthesis, and Beyond. Chem. Biol. 2014, 21 (1), 67–85. 10.1016/j.chembiol.2013.11.011.

(11) Hong, X.; Hopfinger, A. J. Construction, Molecular Modeling, and Simulation of Mycobacterium Tuberculosis Cell Walls. Biomacromolecules 2004, 5 (3), 1052–1065. 10.1021/bm034514c.

(12) Asselineau, J.; Lederer, E. Structure of the Mycolic Acids of Mycobacteria. Nature. 1950, pp 782–783. 10.1038/166782a0.

(13) Watanabe, M.; Aoyagi, Y.; Mitome, H.; Fujita, T.; Naoki, H.; Ridell, M.; Minnikin, D. E. Location of Functional Groups in Mycobacterial Meromycolate Chains; the Recognition of New Structural Principles in Mycolic Acids. Microbiology 2002, 148 (6), 1881–1902. 10.1099/00221287-148-6-1881.

(14) Barkan, D.; Liu, Z.; Sacchettini, J. C.; Glickman, M. S. Mycolic Acid Cyclopropanation Is Essential for Viability, Drug Resistance, and Cell Wall Integrity of Mycobacterium Tuberculosis. Chem. Biol. 2009, 16 (5), 499–509. 10.1016/j.chembiol.2009.04.001.

(15) Dubnau, E.; Chan, J.; Raynaud, C.; Mohan, V. P.; Lanéelle, M. A.; Yu, K.; Quémard, A.; Smith, I.; Daffé, M. Oxygenated Mycolic Acids Are Necessary for Virulence of Mycobacterium Tuberculosis in Mice. Mol. Microbiol. 2000, 36 (3), 630–637. 10.1046/j.1365-2958.2000.01882.x.

(16) Vander Beken, S.; Al Dulayymi, J. R.; Naessens, T.; Koza, G.; Maza-Iglesias, M.; Rowles, R.; Theunissen, C.; De Medts, J.; Lanckacker, E.; Baird, M. S.; Grooten, J. Molecular Structure of the Mycobacterium Tuberculosis Virulence Factor, Mycolic Acid, Determines the Elicited Inflammatory Pattern. Eur. J. Immunol. 2011, 41 (2), 450–460. 10.1002/eji.201040719.

(17) Glickman, M. S.; Cox, J. S.; Jacobs, W. R. A Novel Mycolic Acid Cyclopropane Synthetase Is Required for Cording, Persistence, and Virulence of Mycobacterium Tuberculosis. Mol. Cell 2000, 5 (4), 717–727. 10.1016/S1097-2765(00)80250-6.

(18) Verschoor, J. A.; Baird, M. S.; Grooten, J. Towards Understanding the Functional Diversity of Cell Wall Mycolic Acids of Mycobacterium Tuberculosis. Prog. Lipid Res. 2012, 51 (4), 325–339. 10.1016/j.plipres.2012.05.002.

(19) Barry, C. E.; Lee, R. E.; Mdluli, K.; Sampson, A. E.; Schroeder, B. G.; Slayden, R. A.; Yuan, Y. Mycolic Acids: Structure, Biosynthesis and Physiological Functions. Prog. Lipid Res. 1998, 37 (2–3), 143–179. 10.1016/S0163-7827(98)00008-3.

(20) Radchenko, E. V.; Antonyan, G. V.; Ignatov, S. K.; Palyulin, V. A. Machine Learning Prediction of Mycobacterial Cell Wall Permeability of Drugs and Drug-like Compounds. Molecules 2023, 28 (2). 10.3390/molecules28020633.

(21) Watanabe, M.; Aoyagi, Y.; Ridell, M.; Minnikin, D. E. Separation and Characterization of Individual Mycolic Acids in Representative Mycobacteria. Microbiology 2001, 147 (7), 1825–1837. 10.1099/00221287-147-7-1825.

(22) Ställberg-Stenhagen, S.; Stenhagen, E. A Monolayer and X-Ray Study of Mycolic Acid From the Human Tubercle Bacillus. J. Biol. Chem. 1945, 159 (2), 255–262. 10.1016/s0021-9258(19)52786-7.

(23) Hasegawa, T.; Nishijo, J.; Watanabe, M.; Funayama, K.; Imae, T. Conformational Characterization of α-Mycolic Acid in a Monolayer Film by the Langmuir-Blodgett Technique and Atomic Force Microscopy. Langmuir 2000, 16 (18), 7325–7330. 10.1021/la0004606.

(24) Villeneuve, M.; Kawai, M.; Kanashima, H.; Watanabe, M.; Minnikin, D. E.; Nakahara, H. Temperature Dependence of the Langmuir Monolayer Packing of Mycolic Acids from Mycobacterium Tuberculosis. Biochim. Biophys. Acta - Biomembr. 2005, 1715 (2), 71–80. 10.1016/j.bbamem.2005.07.005.

(25) Zhang, Z.; Pen, Y.; Edyvean, R. G.; Banwart, S. A.; Dalgliesh, R. M.; Geoghegan, M. Adhesive and Conformational Behaviour of Mycolic Acid Monolayers. Biochim. Biophys. Acta - Biomembr. 2010, 1798 (9), 1829–1839. 10.1016/j.bbamem.2010.05.024.

(26) Hasegawa, T.; Amino, S.; Kitamura, S.; Matsumoto, L.; Katada, S. I.; Nishijo, J. Study of the Molecular Conformation of α- and Keto-Mycolic Acid Monolayers by the Langmuir-Blodgett Technique and Fourier Transform Infrared Reflection-Absorption Spectroscopy. Langmuir 2003, 19 (1), 105–109. 10.1021/la026548w.

(27) Hasegawa, T.; Leblanc, R. M. Aggregation Properties of Mycolic Acid Molecules in Monolayer Films: A Comparative Study of Compounds from Various Acid-Fast Bacterial Species. Biochim. Biophys. Acta - Biomembr. 2003, 1617 (1–2), 89–95. 10.1016/j.bbamem.2003.09.008.

(28) Villeneuve, M.; Noguchi, H. Roles of α-Methyl Trans-Cyclopropane Groups in Behavior of Mixed Mycolic Acid Monolayers. Biochim. Biophys. Acta - Biomembr. 2019, 1861 (2), 441–448. 10.1016/j.bbamem.2018.10.019.

(29) Groenewald, W.; Bulacu, M. I.; Croft, A. K.; Marrink, S. J. Molecular Dynamics of Mycolic Acid Monolayers. 2019, 1–46.

(30) Groenewald, W.; Parra-Cruz, R. A.; Jäger, C. M.; Croft, A. K. Revealing Solvent-Dependent Folding Behavior of Mycolic Acids from Mycobacterium Tuberculosis by Advanced Simulation Analysis. J. Mol. Model. 2019, 25 (3). 10.1007/s00894-019-3943-5.

(31) Savintseva, L. A.; Steshin, I. S.; Avdoshin, A. A.; Panteleev, S. V.; Rozhkov, A. V.; Shirokova, E. A.; Livshits, G. D.; Vasyankin, A. V.; Radchenko, E. V.; Ignatov, S. K.; Palyulin, V. A. Conformational Dynamics and Stability of Bilayers Formed by Mycolic Acids from the Mycobacterium Tuberculosis Outer Membrane. Molecules 2023, 28 (3), 1347. 10.3390/molecules28031347.

(32) Esteban-Martín, S.; Jelger Risselada, H.; Salgado, J.; Marrink, S. J. Stability of Asymmetric Lipid Bilayers Assessed by Molecular Dynamics Simulations. J. Am. Chem. Soc. 2009, 131 (42), 15194–15202. 10.1021/ja904450t.

(33) Basu, S.; Mandal, S.; Maiti, P. K. TB Drugs Permeability through Mycolic Acid Monolayer: A Tale of Two Force Fields. Phys. Chem. Chem. Phys. 2024, 19. 10.1039/d4cp02659d.

(34) Ndlandla, F. L.; Ejoh, V.; Stoltz, A. C.; Naicker, B.; Cromarty, A. D.; van Wyngaardt, S.; Khati, M.; Rotherham, L. S.; Lemmer, Y.; Niebuhr, J.; Baumeister, C. R.; Al Dulayymi, J. R.; Swai, H.; Baird, M. S.; Verschoor, J. A. Standardization of Natural Mycolic Acid Antigen Composition and Production for Use in Biomarker Antibody Detection to Diagnose Active Tuberculosis. J. Immunol. Methods 2016, 435, 50–59. 10.1016/j.jim.2016.05.010.

(35) Thanyani, S. T.; Roberts, V.; Siko, D. G. R.; Vrey, P.; Verschoor, J. A. A Novel Application of Affinity Biosensor Technology to Detect Antibodies to Mycolic Acid in Tuberculosis Patients. J. Immunol. Methods 2008, 332 (1–2), 61–72. 10.1016/j.jim.2007.12.009.

(36) Zuber, B.; Chami, M.; Houssin, C.; Dubochet, J.; Griffiths, G.; Daffé, M. Direct Visualization of the Outer Membrane of Mycobacteria and Corynebacteria in Their Native State. J. Bacteriol. 2008, 190 (16), 5672–5680. 10.1128/JB.01919-07.

(37) Minnikin, D. E.; Lee, O. Y.-C.; Wu, H. H. T.; Nataraj, V.; Donoghue, H. D.; Ridell, M.; Watanabe, M.; Alderwick, L.; Bhatt, A.; Besra, G. S. Pathophysiological Implications of Cell Envelope Structure in Mycobacterium Tuberculosis and Related Taxa. In *Tuberculosis - Expanding Knowledge*; Ribon W, Ed.; Intech: London, UK, 2015; pp 145–175. 10.5772/59585.

(38) Plitzko, M.; Engelhardt, H.; Hoffmann, C.; Leis, A.; Niederweis, M. Disclosure of the Mycobacterial Outer Membrane : Cryo-Electron Tomography and Vitreous Sections Reveal the Lipid Bilayer Structure. Proc. Natl. Acad. Sci. U. S. A. 2008, 105 (10), 3963–3967.

(39) Nikaido, H.; Kim, S.-H.; Rosenberg’, E. Y. Physical Organization of Lipids in the Cell Wall of Mycobacterium Chelonae. Mol. Microbiol. 1993, 8 (6), 1025–1030.

(40) Villeneuve, M.; Kawai, M.; Horiuchi, K.; Watanabe, M.; Aoyagi, Y.; Hitotsuyanagi, Y.; Takeya, K.; Gouda, H.; Hirono, S.; Minnikin, D. E. Conformational Folding of Mycobacterial Methoxy- and Ketomycolic Acids Facilitated by α-Methyl Trans-Cyclopropane Groups Rather than Cis-Cyclopropane Units. Microbiol. (United Kingdom) 2013, 159 (PART11), 2405–2415. 10.1099/mic.0.068866-0.

(41) Villeneuve, M.; Kawai, M.; Watanabe, M.; Aoyagi, Y.; Hitotsuyanagi, Y.; Takeya, K.; Gouda, H.; Hirono, S.; Minnikin, D. E.; Nakahara, H. Conformational Behavior of Oxygenated Mycobacterial Mycolic Acids from Mycobacterium Bovis BCG. Biochim. Biophys. Acta - Biomembr. 2007, 1768 (7), 1717–1726. 10.1016/j.bbamem.2007.04.003.

(42) Villeneuve, M.; Kawai, M.; Watanabe, M.; Aoyagi, Y.; Hitotsuyanagi, Y.; Takeya, K.; Gouda, H.; Hirono, S.; Minnikin, D. E.; Nakahara, H. Differential Conformational Behaviors of α-Mycolic Acids in Langmuir Monolayers and Computer Simulations. Chem. Phys. Lipids 2010, 163 (6), 569–579. 10.1016/j.chemphyslip.2010.04.010.

(43) Hanwell, M. D.; Curtis, D. E.; Lonie, D. C.; Vandermeerschd, T.; Zurek, E.; Hutchisoni, G. R. Avogadro: An Advanced Semantic Chemical Editor, Visualization, and Analysis Platform. J. Cheminform. 2012, 4 (1), 17. 10.1186/1758-2946-4-17.

(44) Stroet, M.; Caron, B.; Visscher, K. M.; Geerke, D. P.; Malde, A. K.; Mark, A. E. Automated Topology Builder Version 3.0: Prediction of Solvation Free Enthalpies in Water and Hexane. J. Chem. Theory Comput. 2018, 14 (11), 5834–5845. 10.1021/acs.jctc.8b00768.

(45) Van Der Spoel, D.; Lindahl, E.; Hess, B.; Groenhof, G.; Mark, A. E.; Berendsen, H. J. C. GROMACS: Fast, Flexible, and Free. J. Comput. Chem. 2005, 26 (16), 1701–1718. 10.1002/jcc.20291.

(46) Vega, C.; Abascal, J. L. F. Simulating Water with Rigid Non-Polarizable Models: A General Perspective. Physical Chemistry Chemical Physics. 2011, pp 19663–19688. 10.1039/c1cp22168j.

(47) Berendsen, H. J. C.; Grigera, J. R.; Straatsma, T. P. The Missing Term in Effective Pair Potentials. J. Phys. Chem. 1987, 91 (24), 6269–6271. 10.1021/j100308a038.

(48) Berendsen, H. J. C.; van der Spoel, D.; van Drunen, R. GROMACS: A Message-Passing Parallel Molecular Dynamics Implementation. Comput. Phys. Commun. 1995, 91 (1–3), 43–56. 10.1016/0010-4655(95)00042-E.

(49) MacKerell, A. D.; Bashford, D.; Dunbrack, R. L.; Evanseck, J. D.; Field, M. J.; Fischer, S.; Gao, J.; Guo, H.; Ha, S.; Joseph-McCarthy, D.; Kuchnir, L.; Kuczera, K.; Lau, F. T. K.; Mattos, C.; Michnick, S.; Ngo, T.; Nguyen, D. T.; Prodhom, B.; Reiher, W. E.; Roux, B.; Schlenkrich, M.; Smith, J. C.; Stote, R.; Straub, J.; Watanabe, M.; Wiórkiewicz-Kuczera, J.; Yin, D.; Karplus, M.; Dunbrack, R. L.; Evanseck, J. D.; Field, M. J.; Fischer, S.; Gao, J.; Guo, H.; Ha, S.; Joseph-McCarthy, D.; Kuchnir, L.; Kuczera, K.; Lau, F. T. K.; Mattos, C.; Michnick, S.; Ngo, T.; Nguyen, D. T.; Prodhom, B.; Reiher, W. E.; Roux, B.; Schlenkrich, M.; Smith, J. C.; Stote, R.; Straub, J.; Watanabe, M.; Wiórkiewicz-Kuczera, J.; Yin, D.; Karplus, M. All-Atom Empirical Potential for Molecular Modeling and Dynamics Studies of Proteins. J. Phys. Chem. 1998, 102 (18), 3586–3616. 10.1021/jp973084f.

(50) Jorgensen, W. L.; Maxwell, D. S.; Tirado-Rives, J. Development and Testing of the OLPS All-Atom Force Field on Conformational Energetics and Properties of Organic Liquids. J. Am. Chem. Soc. 1996, 118 (15), 11225–11236. 10.1021/ja9621760.

(51) Wang, J.; Wolf, R. M.; Caldwell, J. W.; Kollman, P. A.; Case, D. A. Development and Testing of a General Amber Force Field. J. Comput. Chem. 2004, 56531 (9), 1157–1174.

(52) Bayly, C. I.; Merz, K. M.; Ferguson, D. M.; Cornell, W. D.; Fox, T.; Caldwell, J. W.; Kollman, P. A.; Cieplak, P.; Gould, I. R.; Spellmeyer, D. C. A Second Generation Force Field for the Simulation of Proteins, Nucleic Acids, and Organic Molecules. J. Am. Chem. Soc. 1995, 117 (19), 5179–5197. 10.1021/ja00124a002.

(53) Oostenbrink, C.; Villa, A.; Mark, A. E.; Van Gunsteren, W. F. A Biomolecular Force Field Based on the Free Enthalpy of Hydration and Solvation: The GROMOS Force-Field Parameter Sets 53A5 and 53A6. J. Comput. Chem. 2004, 25 (13), 1656–1676. 10.1002/jcc.20090.

(54) Essmann, U.; Perera, L.; Berkowitz, M. L.; Darden, T.; Lee, H.; Pedersen, L. G. A Smooth Particle Mesh Ewald Method. J Chem Phys 1995, 103 (1995), 8577–8593. 10.1063/1.470117.

(55) Darden, T.; York, D.; Pedersen, L. Particle Mesh Ewald: An N·log(N) Method for Ewald Sums in Large Systems. J. Chem. Phys. 1993, 98 (12), 10089–10092. 10.1063/1.464397.

(56) Hess, B.; Bekker, H.; Berendsen, H. J. C.; Fraaije, J. G. E. M. LINCS: A Linear Constraint Solver for Molecular Simulations. J. Comput. Chem. 1997, 18 (12), 1463–1472. 10.1002/(SICI)1096-987X(199709)18:12<1463::AID-JCC4>3.0.CO;2-H.

(57) Bussi, G.; Donadio, D.; Parrinello, M. Canonical Sampling through Velocity Rescaling. J. Chem. Phys. 2007, 126 (1), 014101. 10.1063/1.2408420.

(58) Parrinello, M.; Rahman, A. Polymorphic Transitions in Single Crystals: A New Molecular Dynamics Method. J. Appl. Phys. 1981, 52 (12), 7182–7190. 10.1063/1.328693.

(59) Barducci, A.; Bonomi, M.; Parrinello, M. Metadynamics. Wiley Interdiscip. Rev. Comput. Mol. Sci. 2011, 1 (5), 826–843. 10.1002/wcms.31.

(60) Bonomi, M.; Branduardi, D.; Bussi, G.; Camilloni, C.; Provasi, D.; Raiteri, P.; Donadio, D.; Marinelli, F.; Pietrucci, F.; Broglia, R. A.; Parrinello, M. PLUMED: A Portable Plugin for Free-Energy Calculations with Molecular Dynamics. Comput. Phys. Commun. 2009, 180 (10), 1961–1972. 10.1016/j.cpc.2009.05.011.

