## Supplemental Figure S1, Figure S2, Figure S3, Figure S4 for "Self-assembly of mycolic acid in water: monolayer or bilayer"

**Figures**

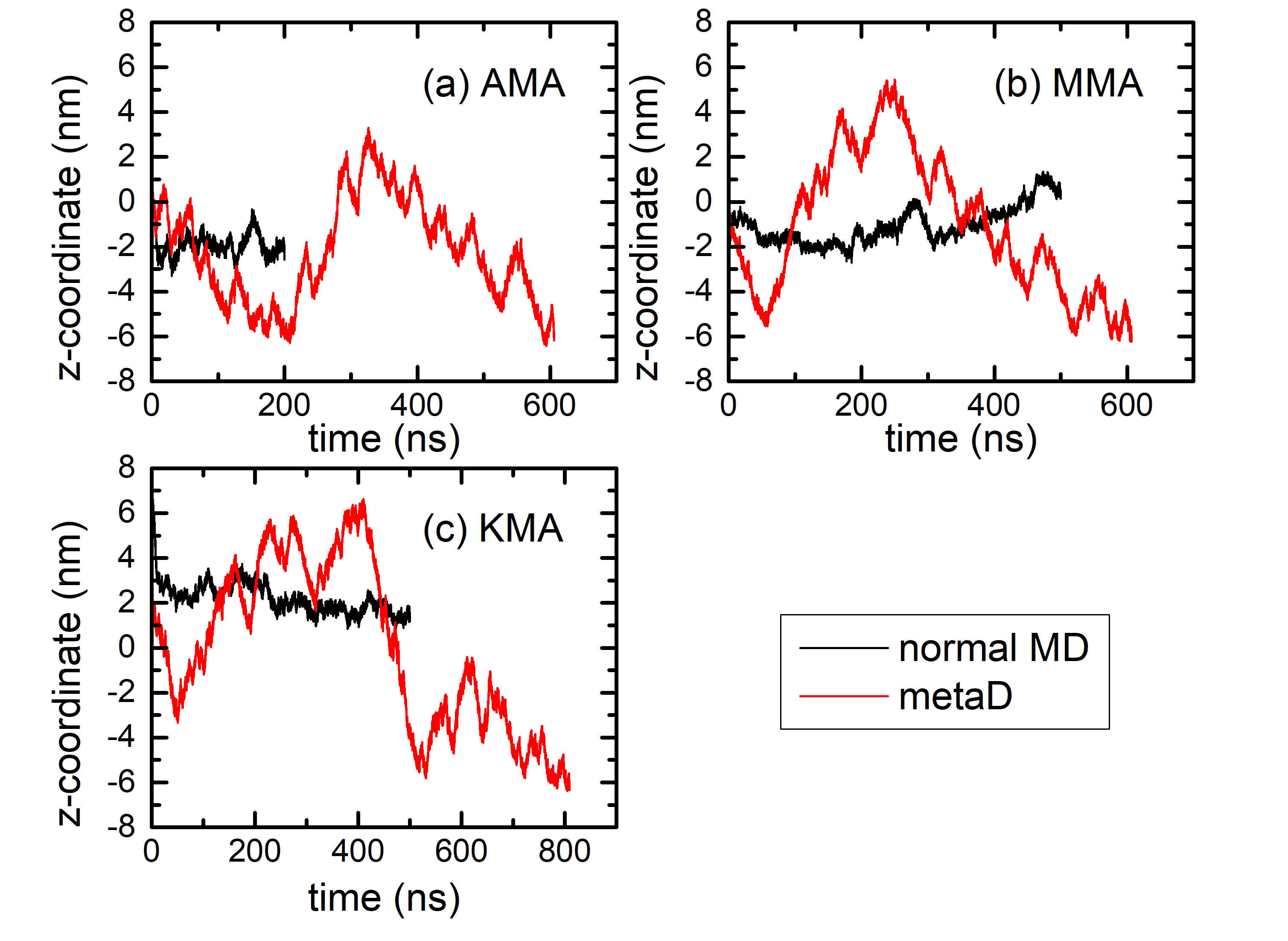

Figure S1. The distance (z-component) of the first two MA chains in the monolayer was estimated through normal MD and metadynamics for (a) AMA, (b) MMA, and (c) KMA in single-component MAs.

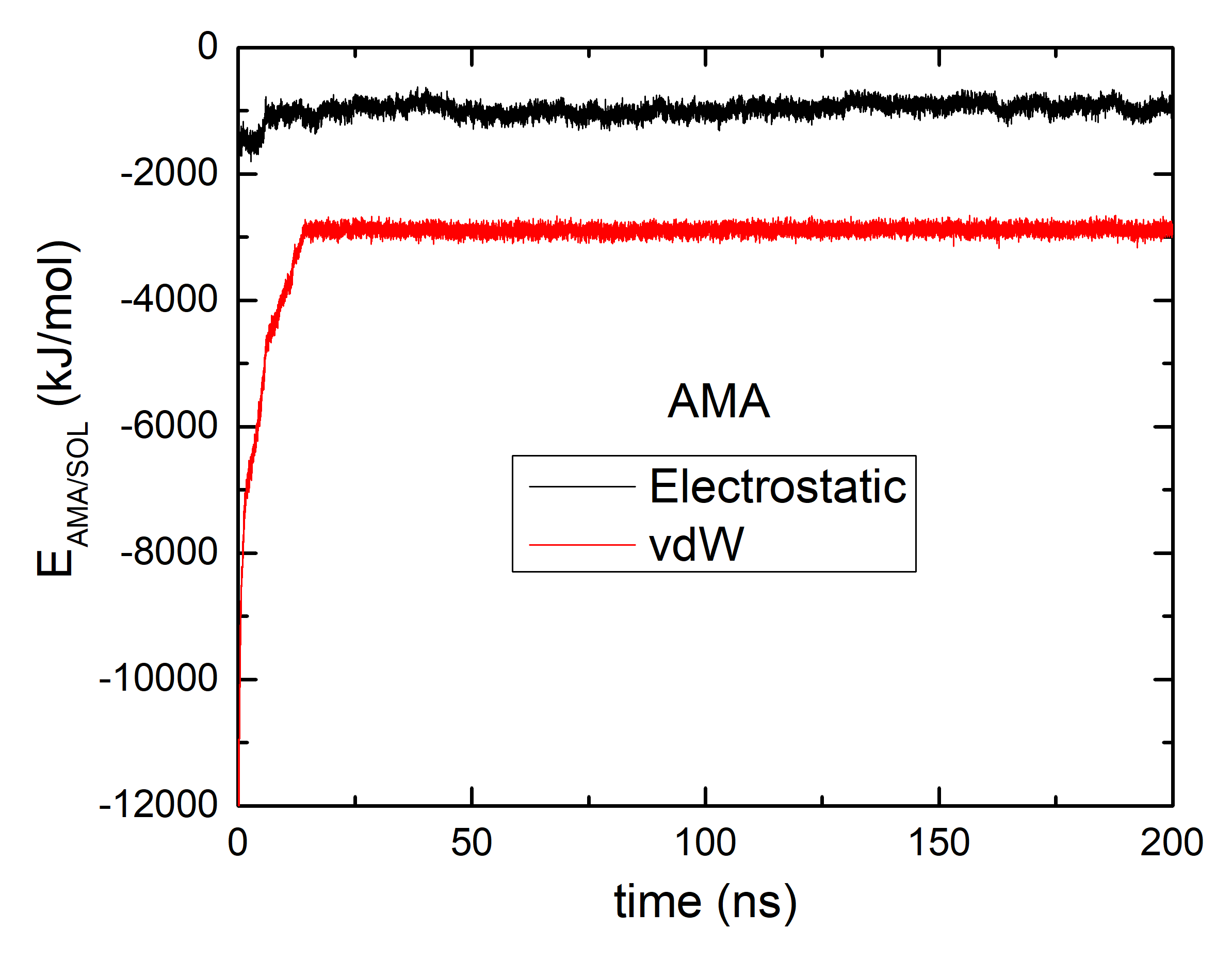

Figure S2. Temporal change in the average non-bonded energy between the alpha-MA (AMA) and solvent molecules.

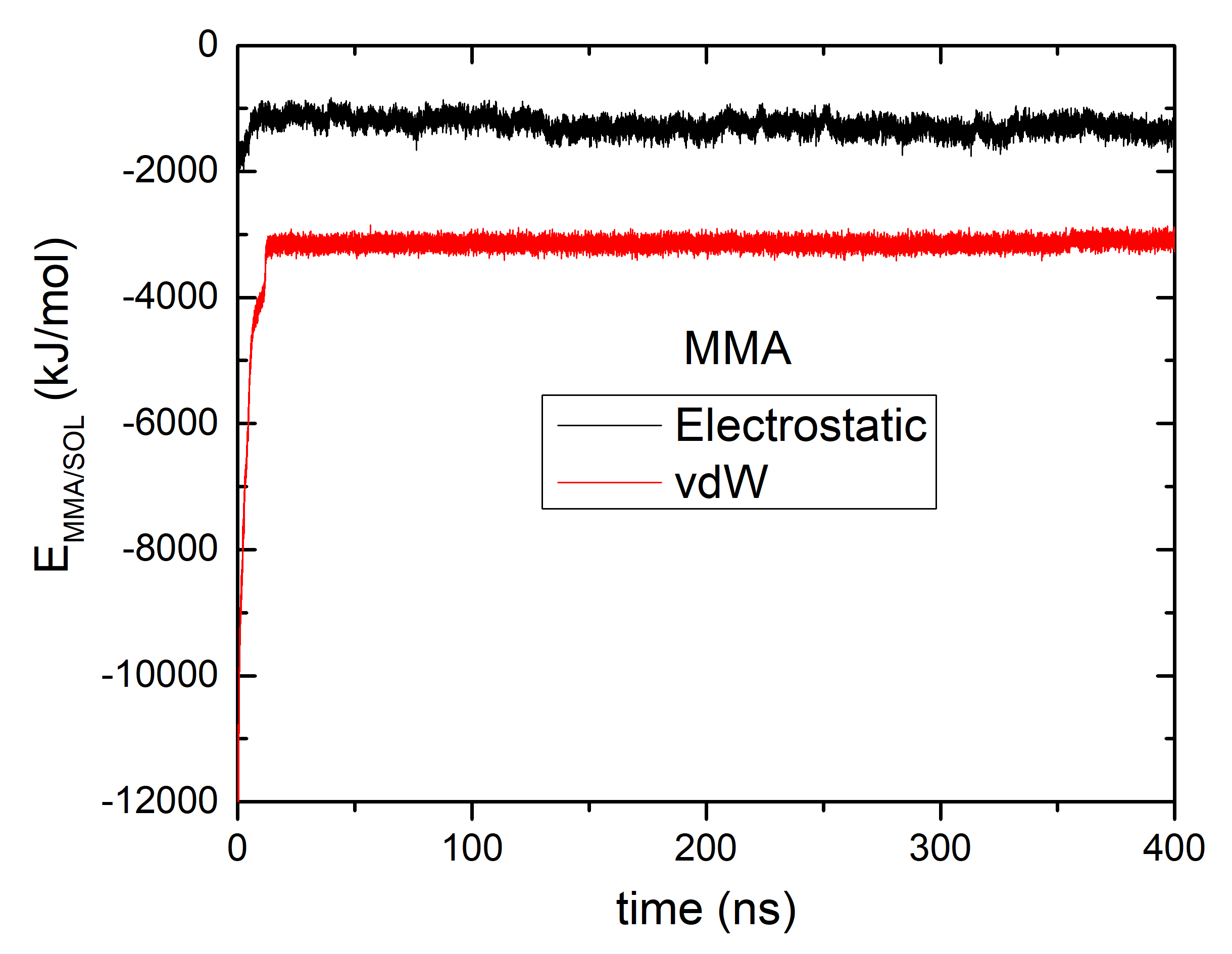

Figure S3. Temporal change in the average non-bonded energy between the methoxy-MA (MMA) and solvent molecules.

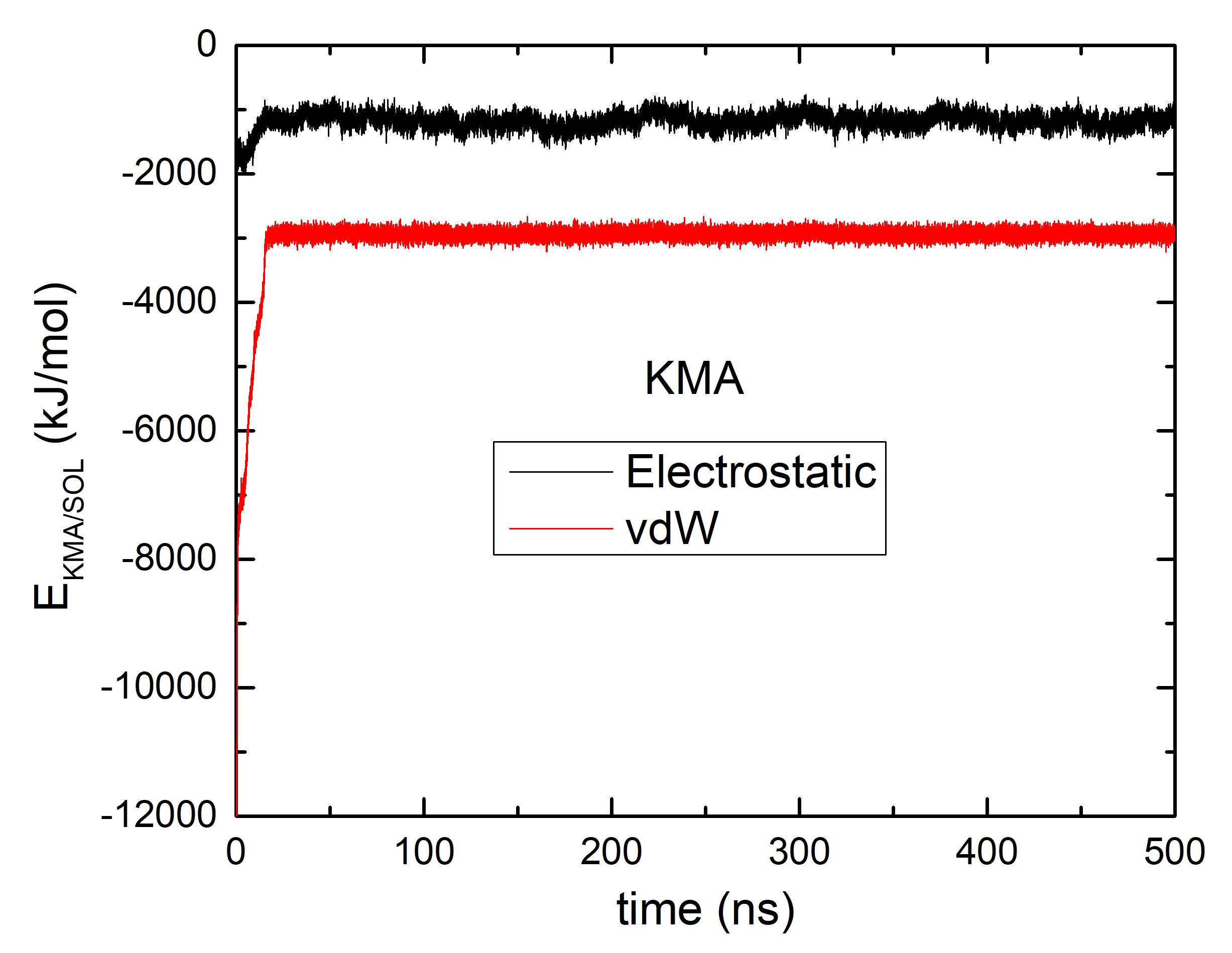

Figure S4. Temporal change in the average non-bonded energy between the keto-MA (KMA) and solvent molecules.
